# SR-BI regulates the synergistic mast cell response by modulating the plasma membrane-associated cholesterol pool

**DOI:** 10.1101/2023.06.21.545859

**Authors:** Sandro Capellmann, Marlies Kauffmann, Michel Arock, Michael Huber

## Abstract

The high-affinity IgE receptor FcεRI is the fundamental mast cell (MC) receptor responsible for the involvement of MCs in IgE-associated allergic disorders. Activation of the FcεRI is achieved via crosslinking by multivalent antigen (Ag) recognized by IgE, which results in degranulation and pro-inflammatory cytokine production. In comparison to the T and B cell receptor complexes, for which several co-receptors that orchestrate the initial signaling have been described, information is scarce about FcεRI-associated proteins. Additionally, it is not completely clear how FcεRI signaling synergizes with input from other receptors and how potential regulators affect this synergistic response. We aimed at identifying new regulators of FcεRI and found that the HDL receptor SR-BI (gene name: *Scarb1*/*SCARB1*) is expressed in MCs, functionally associates with FcεRI and regulates the local plasma membrane cholesterol content in cholesterol-rich plasma membrane nanodomains as shown by using the cholesterol-sensitive probe GFP-D4. This impacted on the activation of murine MCs upon co-stimulation of the FcεRI with different receptors known to synergize with FcεRI-signaling pathways. Amongst them we investigated the co-activation of the FcεRI with the receptor tyrosine kinase KIT, the IL-33 receptor and GPCRs activated by adenosine or PGE_2_. *Scarb1*-deficient bone marrow-derived MCs (BMMCs) showed reduced cytokine secretion in response to these co-stimulation conditions suggesting a role for plasma membrane-associated cholesterol regulating MC-driven inflammation. Mimicking *Scarb1* deficiency by membrane cholesterol depletion employing MβCD, we identified PKB and PLCγ1 as cholesterol-sensitive signaling molecules activated downstream of FcεRI in BMMCs. Specifically, when murine MCs were stimulated with SCF and Ag in combination, PLCγ1 activation appeared to be drastically boosted and this could be mitigated by cholesterol depletion. Inhibiting SR-BI in BMMCs phenocopied this effect. Similarly, SR-BI inhibition also attenuated the synergistic response to PGE_2_ and anti-IgE in the human ROSA^KIT^ ^WT^ mast cell line suggesting that SR-BI is a crucial regulator of synergistic MC activation by regulating the local plasma membrane cholesterol concentration.

## Introduction

The high-affinity IgE receptor FcεRI is the fundamental mast cell (MC) receptor responsible for the involvement of MCs in IgE-associated allergic disorders (1). Activation of the FcεRI is achieved via crosslinking by multivalent allergen/antigen (Ag) recognized by IgE (2). This Ag-dependent FcεRI aggregation results in an immediate release of pre-formed mediators including proteases, biogenic amines and proteoglycans from cytoplasmic granules (3). Additionally, the *de novo* synthesis and secretion of arachidonic acid metabolites and pro-inflammatory cytokines is initiated. The initial activation of the FcεRI is mediated by phosphorylation of the immunoreceptor tyrosine-based activation motifs (ITAMs) in the β- and γ-subunits by the SRC family kinase (SFK) LYN providing docking sites for downstream signaling proteins containing SRC-homology 2 (SH2) domains (4,5). Interestingly, although the phosphorylation of the FcεRI β-subunit ITAMS by LYN is considered the first step in FcεRI activation, how this process is regulated is poorly understood. It is assumed that probably the plasma membrane (PM) structure, notably lipid rafts, and/or steric effects like the displacement of negative regulatory phosphatases or the spatial proximity of multiple FcεRI monomers induced by Ag-dependent crosslinking control ITAM phosphorylation (6). In comparison to other multichain immune recognition receptors (MIRRs) like the T and B cell receptor complexes, for which several co-receptors orchestrating the initial signaling have been described (7,8), information is scarce about FcεRI-associated proteins regulating the early activation steps.

One central signaling enzyme downstream of LYN enabling MC degranulation and cytokine production is phospholipase C gamma 1 (PLCγ1). PLCγ1 hydrolyzes phosphatidyl-inositol-(4,5)-bisphosphate (PI-4,5-P_2_) to diacylglycerol (DAG) and inositol-1,4,5-trisphosphate (IP_3_). This induces an Ag-dependent store-operated calcium (Ca^2+^) entry (SOCE) and protein kinase C (PKC) activation leading to fusion of the cytoplasmic granules with the PM releasing granule-stored mediators (9,10). Besides, Ca^2+^ mediates the activation of the phosphatase calcineurin, which activates the transcription factor NFAT (nuclear factor of activated T cells) essential for transcription of pro-inflammatory cytokines (11). A second crucial pathway in FcεRI signaling involves the LYN-independent, FYN-regulated activation of phosphoinositide-3-kinase (PI3K), which catalyzes the generation of phosphatidylinositol-(3,4,5)-trisphosphate (PI-3,4,5-P_3_) from PI-4,5-P_2_. Importance of the PI3K pathway for FcεRI activation stems from the observation that murine bone marrow-derived MCs (BMMCs) deficient for the negative regulator of the PI3K pathway SH2 domain-containing inositol-5’-phosphatase 1 (SHIP1) show a hyperactive phenotype of enhanced degranulation and pro-inflammatory cytokine production in response to Ag stimulation (12). PI-3,4,5-P_3_ enables membrane recruitment of pleckstrin homology (PH) domain-containing proteins like protein kinase B (PKB/AKT), PDK1 (3-phosphoinositide-dependent kinase 1) or Bruton’s tyrosine kinase (BTK) (13). PKB is activated by PDK1 (14) and regulates the Ag-elicited cytokine response by phosphorylating IκBα, the inhibitor of NF-κB, thus releasing the transcription factor NF-κB from its inhibitory complex and activating NF-κB-dependent transcription (15).

Apart from the FcεRI, MCs possess a plethora of other receptors to adequately respond to pathogens or tissue barrier disruption as they are strategically located at host-environment interfaces (1). Some of these receptors have been shown to boost the reaction of MCs to Ag when minute amounts of Ag would only elicit a poor response. For instance, co-activation of the receptor tyrosine kinase (RTK) KIT by stem cell factor (SCF) and the FcεRI by Ag leads to a synergistic enhancement of degranulation and pro-inflammatory cytokine production, which implies that the signaling pathways of both receptors are interconnected. The synergism of SCF and Ag has been shown to either depend on PI3K/PKB and/or on PKCβ activation (16,17). Likewise, synergistic responses affecting degranulation and/or cytokine production of MCs have also been shown to occur upon co-stimulation of Ag with IL-33 (18), Toll-like receptor (TLR) 2 and TLR4 agonists (19), and GPCR agonists like adenosine (A_3_ receptor) and PGE_2_ (EP3 receptor) (20,21). Nevertheless, the question addressing the underlying mechanisms involving the synergistic responses is hard to answer as relevant signaling pathways are often activated by both stimuli, making it difficult to assign context-dependent functions of one signaling element to one or the other receptor.

Scavenger receptor class B member I (SR-BI, gene name: *SCARB1*/*Scarb1*) is the canonical receptor for high-density lipoprotein (HDL) (22). SR-BI is the central player in reverse cholesterol transport and mainly expressed in the liver and steroidogenic organs to clear the body from excess cholesterol or to supply cholesterol for steroid hormone synthesis, respectively (23). Besides, SR-BI is also expressed in macrophages and endothelial cells where it has positive as well as negative regulatory functions in atherosclerosis. Macrophage SR-BI mediates efferocytosis while endothelial SR-BI promotes the transport of low-density lipoprotein (LDL) into the artery wall (24,25). While the former is attenuates atherosclerotic progression, the latter fosters atherosclerosis development. Unlike the LDL receptor, SR-BI mediates the bi-directional transport of HDL lipids and especially cholesterol between the HDL particle and the PM *via* a shuttling mechanism avoiding internalization of the complete lipoprotein particle, thereby regulating the cellular and especially the PM cholesterol homeostasis (26–28). Of note, the organization of the PM in cholesterol-rich lipid-ordered (also known as lipid rafts) and lipid-disordered regions has been shown to contribute to signaling regulation. Lipid rafts are commonly related to organization of signaling at the PM providing platforms of protein complex assembly (29). This has led to the general view that lipid rafts are important for proper signaling initiation downstream of MIRRs, RTKs and GPCRs (30–33). However, this model has been challenged by observations that cholesterol can also have a negative regulatory role for receptor activation. For instance, the TCR β chain can bind cholesterol, which keeps the receptor in an inactive, resting state (34). Besides, mild lipid raft disruption by methyl-β-cyclodextrin (MβCD) in T cells was sufficient to trigger and augment anti-CD3-induced tyrosine phosphorylation (35). Thus, the exact role of cholesterol and lipid rafts is still unclear and may strongly depend on the cellular state and cell type analyzed.

In this study we show that SR-BI is expressed in different MC models, functionally associates with the FcεRI in murine MCs and regulates the synergistic enhancement of cytokine production in response to co-activation of the FcεRI with KIT, ST2 or GPCRs. *Scarb1^-/-^* BMMCs displayed reduced focally enriched cholesterol in the PM, which we could mimic using MβCD or cholesterol oxidase. We could show that the PKB and PLCγ1 pathways activated by Ag stimulation are both cholesterol-dependent. MβCD-mediated cholesterol extraction strongly attenuated PKB and PLCγ1 activation. Besides, cholesterol oxidation and SR-BI inhibition in bone marrow-derived MCs (BMMCs) specifically reduced PLCγ1 signaling upon co-stimulation of KIT and FcεRI without affecting signaling downstream of individual receptor activation. These data provide a model for SR-BI’s role in determining PM homeostasis and regulating synergistic MC activation.

## Material and Methods

### Animals and Cell Culture

All mice used in this study were on a mixed C57BL/6 x 129/Sv background. Experiments were performed using *Scarb1^-/-^* transgenic (36) and corresponding wildtype (WT) mice. Murine bone marrow cells were isolated and BMMCs were differentiated as described previously (37). BMMCs were cultivated as single cell suspensions in growth medium (RPMI 1640 medium (Invitrogen, #21875-0991) containing 15% FCS (Capricorn, #FBS-12A), 10 mM HEPES, 50 units/ml Penicillin and 50 mg/ml Streptomycin, 100 µM β-mercaptoethanol, 30 ng/ml interleukin 3 (IL-3) from X63-Ag8-653 conditioned medium (38)). PMCs were cultivated in the same way but additionally supplemented with 20 ng/ml SCF derived from cell culture supernatant from CHO cells that were transfected with an expression vector expressing murine SCF (37). The differentiation of BMMCs and the enrichment of PMCs were evaluated after 4-5 weeks in culture by FACS analysis (see below) and was considered successful if >95% of BMMCs or PMCs were positive for MC surface markers FcεRI, KIT, ST2 and CD13. Experiments were performed in accordance with German legislation governing animal studies and following the principles of laboratory animal care. Mice are held in the Institute of Laboratory Animal Science, Medical Faculty of RWTH Aachen University. The institute holds a license for husbandry and breeding of laboratory animals from the veterinary office of the Städteregion Aachen (administrative district). The institute follows a quality management system, which is certified according to DIN EN ISO 9001:2015. Every step in this project involving mice was reviewed by the animal welfare officer. All experiments were approved by the Landesamt für Natur, Umwelt, und Verbraucherschutz NRW (LANUV), Recklinghausen.

HMC-1.1 and −1.2 cell lines were cultivated in RPMI1640 medium (Invitrogen, #21875-0991) containing 10% FCS (Capricorn, #FBS-12A). ROSA^KIT WT^ cells were cultivated according to (39) without addition of IL-6 or IL-3. Medium was supplemented with murine SCF from CHO cell culture supernatant. ROSA^KIT D816V^ cells were grown in IMDM medium (Gibco, #11504556) containing 10% FCS (Capricorn, #FBS-12A).

### β-Hexosaminidase Assay

To measure degranulation, MCs were pre-loaded with 0.15 µg/ml IgE (clone Spe-7, Sigma, #A2831) overnight (37°C, 5% CO_2_). Cells were washed in PBS and concentration was adjusted to 1.2 x 10^6^ cells/ml in Tyrode’s buffer (130 mM NaCl, 5 mM KCl, 1.4 mM CaCl_2_, 1 mM MgCl_2_, 5.6 mM glucose, and 0.1% BSA in 10 mM HEPES, pH 7.4). After adaptation to 37°C for 15 min, cells were stimulated as indicated with different concentrations of Ag (DNP-HSA, Sigma, #A6661). The degree of degranulation was determined by quantification of β-hexosaminidase release as described previously (40).

### Pro-inflammatory Cytokine ELISA

To determine IL-6 and TNF secretion, MCs were stimulated as indicated in the respective experiments. If cells were stimulated with Ag, MCs were pre-loaded with IgE (clone Spe-7, 0.15 µg/ml). Cell number was adjusted to 1.2 x 10^6^ cells/ml in stimulation medium (RPMI 1640 + 0.1% BSA (Serva, #11930.04)), cells were allowed to adapt to 37°C and stimulated for 3 hours. Supernatants were collected to determine cytokine release. 96-well ELISA plates (Corning, #9018) were coated with capturing anti-IL-6 (1:250, BD Biosciences, #554400) or anti-TNF (1:200, R&D Systems, #AF410-NA) antibodies diluted in PBS overnight at 4°C according to manufacturer’s instructions. ELISA plates were washed three times with PBS+0.1% Tween and blocked with PBS+2% BSA (IL-6 ELISA) or PBS+1%BSA+5% sucrose (TNF ELISA) before loading of supernatants (50 µl for IL-6 ELISA, 100 µl for TNF ELISA). Additionally to loading of supernatants, IL-6 (BD Pharmingen, #554582) and TNF (R&D Systems, #410-MT-010) serial 1:2 standard dilutions were added and plates were incubated overnight at 4°C. Thereupon, plates were washed three times again followed by incubation with biotinylated anti-IL-6 (1:500, BD Biosciences, #554402) and anti-TNF (1:250, R&D Systems, #BAF-410) antibodies diluted in PBS+1% BSA for 45 minutes and 2 hours, respectively, at room temperature (RT). After 3 washing steps, streptavidin alkaline phosphatase (SAP, 1:1000, BD Pharmingen, #554065) was added for 30 minutes at RT. After 3 more washing steps, the substrate *p*-Nitro-phenyl-phosphate (1 pill per 5 ml in sodium carbonate buffer (2mM MgCl_2_ in 50 mM sodium carbonate, pH 9.8), Sigma, #S0942-200TAB) was added and OD_450_ was recorded using a plate reader (BioTek Eon). Qualitative differences and similarities between WT and mutant cells were consistent throughout the study. Levels of secreted cytokines varied due to batch-to-batch variations of primary, differentiated cells of different age and from different mice.

### Flow Cytometry, LAMP1 Assay, Co-internalization Assay and Plasma Membrane GM1 and Cholesterol Staining

To confirm the MC identity of individual, *in vitro* differentiated BMMC cultures, expression of FcεRI, KIT, ST2 and CD13 was verified using FITC-coupled anti-FcεRI (1:100, clone MAR-1, BioLegend, #134306), PE-coupled anti-KIT (1:100, clone 2B8, BioLegend, #105808), FITC-coupled anti-ST2 (1:100, clone DJ8, MD Bioproducts, #101001F) and PE-coupled anti-CD13 (1:100, clone R3-242, BD Pharmingen, #558745) antibodies. Roughly 0.5 x 10^6^ cells were washed in FACS buffer (PBS containing 3% FCS) and stained with the indicated antibodies for 20 minutes at 4°C in the dark. Thereupon, cells were washed again in FACS buffer and analyzed by flow cytometry. SR-BI cell surface expression was determined accordingly using anti-SR-BI antibody (1:100, USBiological, #S0099-01C) and PE-coupled anti-rabbit secondary antibody (1:200, Dianova, # 711-116-152). Before antibody incubation, cells were incubated with Fc-block (clone 2.4G2, BD Biosciences, #553141) for 10 min at RT.

For LAMP1 assay, IgE preloaded BMMCs were washed in PBS and concentration was adjusted to 1 x 10^6^ cells/ml in stimulation medium. Cells were stimulated with indicated Ag concentrations and for indicated time. Reaction was stopped by centrifugation at 4°C and externalized LAMP1 (CD107a) was stained using FITC-coupled anti-LAMP1 antibody (clone 1D4B, BioLegend, #121605) for 20 minutes at 4°C in the dark. Subsequently, cells were washed in FACS buffer and analyzed by flow cytometry.

Co-internalization of FcεRI and SR-BI was analyzed upon Ag stimulation of BMMCs similar to LAMP1 assay. Cell surface localization of FcεRI and SR-BI was determined by using FITC-labelled anti-FcεRI antibody (Clone MAR-1) and anti-SR-BI (USBiological) and PE-labelled anti-rabbit antibodies (Dianova) following blocking of FcγRs with Fc-block (BD Biosciences).

Staining of PM high-cholesterol regions was performed using the His- and GFP-tagged D4 domain of *Clostridium perfringens* exotoxin (10 µg/ml) as described previously (41). The plasmid encoding His-tagged GFP-D4 was provided by RIKEN Institute (RDB13961 clone: pET28/His6-EGFP-D4) deposited by Shimada et al. (42). Ganglioside M1 (GM1) staining was performed using recombinant Choleratoxin B labelled with AlexaFluor647 (3 µg/ml, Thermo Fisher, #C34778). Both GFP-D4 and AlexaFluor647-labelled CTB were added to FACS buffer-washed cells and incubated for 20 minutes at 4°C in the dark. Cells were washed again before analysis by flow cytometry.

All flow cytometry analyses described herein were performed using a FACSCanto II (BD Biosciences). Acquired data were analyzed using FlowJo software (Tree Star, Ashland, OR, USA).

### Recombinant protein Purification

His-tagged GFP-D4 was purified from *Escherichia coli* BL21 strain as described in (43).

### RT-qPCR

Total RNA was isolated from 3 x 10^6^ cells using NucleoSpin RNA Plus Kit (Macherey Nagel, #740955.50) according to manufacturer’s instructions. An amount of 1 µg of RNA was reverse transcribed using Omniscript Reverse Transcription (RT) Kit (Qiagen, #205113) together with random oligonucleotides (Roche, #11034731.001) according to manufacturer’s instructions. Transcript expression was quantified using Sybr green reaction mix SensiFAST (Bioline, #BIO-86005) and 10 pmol of specific primers. PCR reaction was performed on a Rotorgene Q (Qiagen). Expression of *Hprt/HPRT* served as housekeeping gene to determine relative expression according to the delta-delta-C_T_ or delta-C_T_ method (44). The following murine primers were used: *Hprt* fwd GCTGGTGAAAAGGACCTCC, *Hprt* rev CACAGGACTAGAACACCTGC; *Il6* fwd TCCAGTTGCCTTCTTGGGAC, *Il6* rev GTGTAATTAAGCCTCCGACTTG; *Tnf* fwd AGCACAGAAAGCATGATCCGC, *Tnf* rev TGCCACAAGCAGGAATGAGAAG; *Ptgs2* fwd ATCCCCCCACAGTCAAAGACAC, *Ptgs2* rev ACATCATCAGACCAGGCACCAG; *Scarb1* fwd: GTCATGATCCTCATGGTGCC, *Scarb1* rev TTCGAAG AAGTAGACAGACAAGT. The following human primers were used: *HPRT* fwd TGACACTGGCAAAACAATGCA, *HPRT* rev GGTCCTTTTCACCAGCAAGCT; *SCARB1* fwd TGCTCTGGTTTGCAGAGAGC, *SCARB1* rev TTTGGCAGAT GACAGGGACC; *IL8* fwd CACTGCGCCAACACACAGAAAT, *IL8* rev ATGAATT CTCAGCCCTCTTCAA; *TNF* fwd ACTTTGGAGTGATCGGGCC, *TNF* rev CATTGGCCAGGAGGGCATT; *CCL1* fwd TGCGGAGCAAGAGATTCCC, *CCL1* rev ACCCATCCAACTGTGTCCAAG.

### Stimulation of MCs and Western Blotting

To analyze signaling by Western blot, BMMCs were starved overnight (RPMI 1640+10% FCS without IL-3) and IgE-pre-loaded. For stimulation, cell number was adjusted to 1 x 10^6^/ml in stimulation medium. ROSA^KIT WT^ cells were seeded at 0.3 x 10^6^ cells/ml and treated for 4 days with IL-4 (20 ng/ml, PeproTech, #200-04) and human IgE (1 µg/ml, Sigma, #401152) in regular culture medium according to (39) before stimulation. Cells were treated or stimulated as indicated in the respective experiments and snap-frozen in liquid nitrogen to stop the reaction. 1 x 10^6^ cells were lysed in phosphorylation solubilization (PSB) buffer (50 mM HEPES, 100 mM sodium fluoride, 10 mM sodium pyrophosphate, 2 mM sodium orthovanadate, 2 mM EDTA, 2 mM sodium molybdate, 0.5% NP-40, 0.3% sodium deoxycholate and 0.03% sodium dodecylsulfate (SDS)) for 30 minutes at 4°C. Lysates were centrifuged (10 minutes, 130000 x g) and analyzed by SDS-PAGE and Western blot as described previously (45).

### Cellular Cholesterol Content Determination

To determine total (unesterified and esterified) cellular cholesterol, Amplex Red Cholesterol Kit (Thermo Fisher, #A12216) was used according to the manufacturer’s instructions.

### Statistical Analysis

Presented data were generated from at least three independent experiments. Graphical illustration and statistical analyses of data were performed using GraphPad Prism 9 (GraphPad Software, San Diego, CA 92108). Statistical test procedures were performed as indicated in the respective figure legends. *P* values were considered statistically significant according to the GraphPad style (ns: *p*>0.05, * *p*<0.05, ** *p*<0.01, *** *p*<0.001, **** *p*<0.0001). The respective number of independent biological replicates per experiment is indicated in the corresponding figure legends.

## Results

### Ezetimibe inhibits MC effector functions independent of CD13

Ezetimibe is a licensed LDL-derived cholesterol-lowering drug often administered in combination with statins to reduce the risk of cardiovascular events (46). Ezetimibe was reported to have multiple targets to which it can bind and potentially exert its effects as a cholesterol absorption inhibitor. One of the reported interactors was CD13 (47). Coincidentally, CD13 is expressed in MCs and is reported to be a negative regulator of Ag-triggered MC activation (48). Therefore, we asked if Ezetimibe is able to mimic the *Cd13^-/-^* phenotype of enhanced degranulation and cytokine production upon FcεRI activation (48). Surprisingly, Ezetimibe dose-dependently inhibited Ag-elicited degranulation and IL-6 production in wild type (WT) BMMCs (Fig. 1A & B). Moreover, in *Cd13^-/-^* BMMCs Ezetimibe inhibited degranulation (Fig. 1C), suggesting that additional molecular targets for Ezetimibe exist apart from CD13 in MCs. We evaluated, which signaling pathways in MCs are affected by Ezetimibe. We pre-treated BMMCs for one hour with different concentrations of Ezetimibe and subsequently stimulated the FcεRI with an optimal dose of Ag (20 ng/ml). In the DMSO control-treated WT BMMCs, we could detect induction of phosphorylation of threonine 308 (T308) and serine 473 (S473) of PKB, as well as activation of the mitogen-activated protein kinases (MAPKs) p38 and ERK1/2 (Fig. 1D). Ezetimibe treatment predominantly inhibited initial PKB T308 and at later time points PKB S473 phosphorylation, while ERK1/2 and p38 activation were only slightly diminished at later time points (Fig. 1D & E). Collectively, these data point to a CD13-independent inhibitory function of Ezetimibe on FcεRI-dependent MC activation affecting predominantly activation of the PI3K-PKB pathway.

**Fig. 1:**
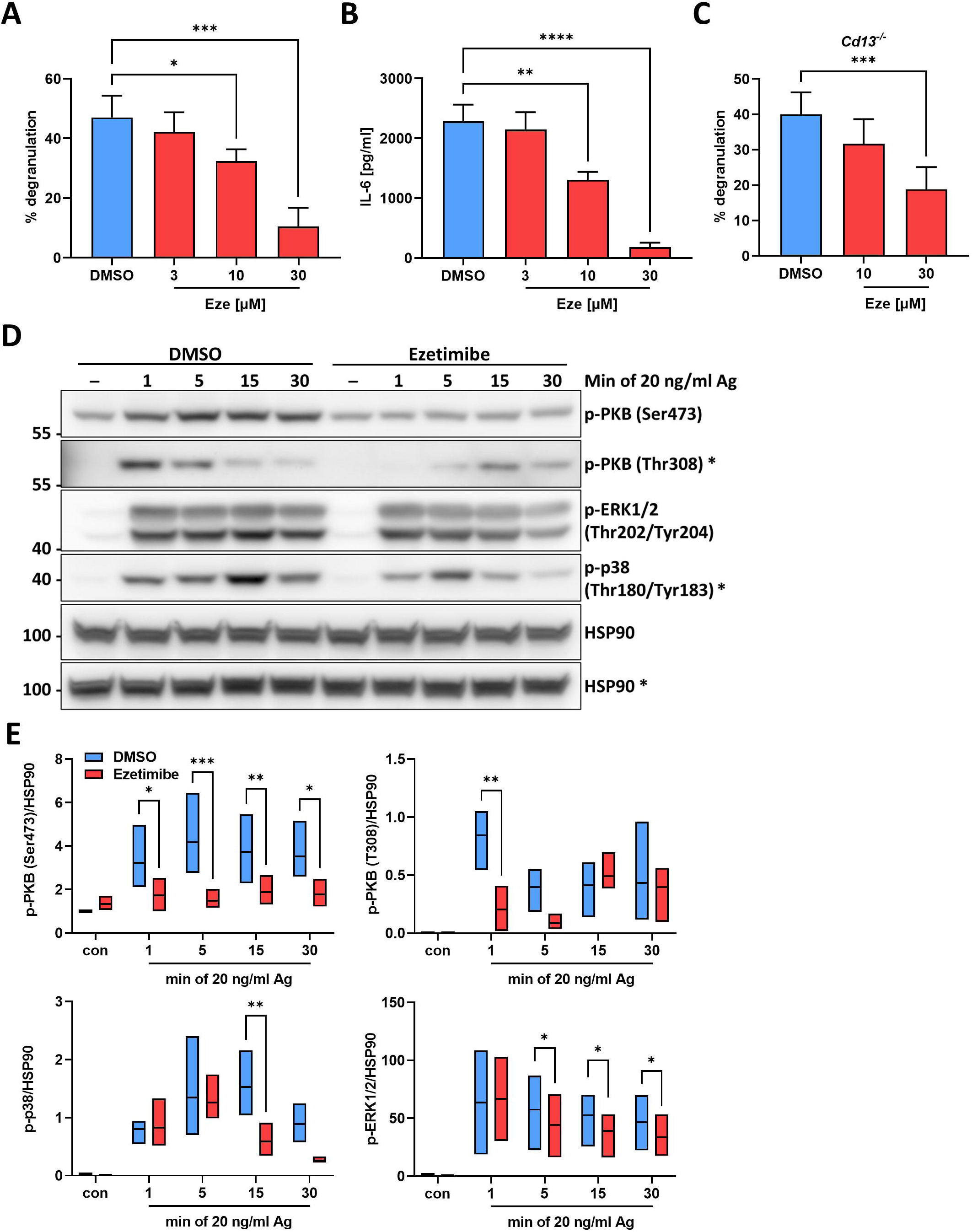
Ezetimibe inhibits mast cell effector functions independent of CD13. **(A)** WT BMMCs were pre-treated with indicated concentrations of Ezetimibe for 1 hour and subsequently stimulated with Ag [20 ng/ml] for 30 minutes The extent of degranulation was determined by quantification of released β-hexosaminidase (n=3). **(B)** WT BMMCs were treated with Ezetimibe and stimulated as in **(A)**. The amount of secreted IL-6 was determined by ELISA after 3 hours of stimulation (n=3). **(C)** *Cd13^-/-^* BMMCs were pre-treated with indicated Ezetimibe concentrations for 1 hour and then stimulated with 20 ng/ml Ag for 30 minutes. The degranulation was determined by β-hexosaminidase assay (n=3). **(D)** WT BMMCs were pre-treated with DMSO or 30 µM Ezetimibe for 1 hour and subsequently stimulated with 20 ng/ml Ag for the indicated time. Activation of PKB and MAPK signaling pathways p38 and ERK was determined by phospho-specific antibodies in Western blot. HSP90 served as loading control. **(E)** Quantification of Western blot results shown in **(D)**. Activation of signaling pathways was determined by densitometry analysis quantifying the phosphorylated protein in relation to HSP90 (n=3). Data are shown mean +SD. Floating bars indicate range between the minimal and maximal values of the independent replicates, the line indicates the mean value. **(A)**, **(B)** and **(C)** were analyzed by ordinary one-way ANOVA followed by Dunnett test to correct for multiple comparisons. **(E)** was analyzed by two-way ANOVA followed by Sídák multiple comparisons test. * *p*<0.05, ** *p*<0.01, *** *p*<0.001, **** *p*<0.0001.

### SR-BI is expressed in different MC types and can be targeted by BLT-1 revealing a role in FcεRI-mediated MC activation

Apart from CD13, several other possible proteins targeted by Ezetimibe have been reported and have led to partially conflicting and contradictory results regarding the actual mechanism behind the cholesterol-lowering effect of Ezetimibe. These proteins include SR-BI, CD36 and Niemann-Pick C1-like 1 (NPC1L1) (49–53). We analyzed existing transcriptomic data of resting BMMCs and determined basal mRNA expression levels of *Scarb1*, *Cd36* and *Npc1l1* using *Cd13*, *Ms4a2* encoding the β-chain of the FcεRI and *Cd79a* encoding the BCR α-chain as positive and negative controls, respectively. Aside from CD13, the only additional Ezetimibe target found to be expressed in BMMCs on mRNA level above the threshold was *Scarb1* (Fig. 2A). To confirm this, we measured *Scarb1* expression in the two murine MC models, BMMCs and peritoneum-derived MCs (PMCs) as well as in the human MC lines HMC-1.1, HMC-1.2, ROSA^KIT WT^ and ROSA^KIT D816V^. *Scarb1* was higher expressed in BMMCs compared to PMCs, while both HMC cell lines and ROSA^KIT WT^ cells showed a comparable *SCARB1* expression. Interestingly, in ROSA^KIT D816V^ cells *SCARB1* expression was significantly increased compared to the respective WT cells (Fig. 2B). The same relative expression differences were also observed on protein level assessing SR-BI expression using an antibody specifically recognizing both murine and human SR-BI (Fig. 2C & D). Using *Scarb1^-/-^* BMMCs, we addressed the role of SR-BI for the effect of Ezetimibe on degranulation and cytokine production upon FcεRI stimulation. Intriguingly, *Scarb1^-/-^* BMMCs were still sensitive to Ezetimibe-mediated inhibition of degranulation and cytokine production (suppl. Fig. 1A & B) pointing to further off-target effects on other cellular targets. However, we also tested the SR-BI selective inhibitor BLT-1 (blocker of lipid transfer 1) in the context of Ag-triggered MC activation to reveal if SR-BI modulates FcεRI activation. Although pre-treatment of BMMCs with BLT-1 had no effect on degranulation, which we quantified by determination of β-hexosaminidase release (suppl. Fig. 1C), we could detect a slightly but consistently reduced production and secretion of IL-6 and TNF in response to an optimal concentration of Ag and IL-33 stimulation, while the responses to SCF, LPS and FSL-1 were unaffected by SR-BI inhibition (Fig. 2E). Interestingly, similar to Ezetimibe, BLT-1 reduced the activation of PKB at later time points (15 min) in WT BMMCs upon sub-optimal Ag stimulation. Besides, activation of PLCγ1 and the MAPK p38 were reduced after 15 min of sub-optimal Ag stimulation, while phosphorylation of ERK1 and ERK2 was not altered. Similarly, phosphorylation signals of all assessed proteins were unaltered after 5 min of Ag stimulation (suppl. Fig. 1D). Hence, we concluded that SR-BI has no impact on the immediate FcεRI-dependent MC activation (up to 5 min of Ag stimulation) but may regulate later stages of pro-inflammatory MC responses.

**Fig. 2:**
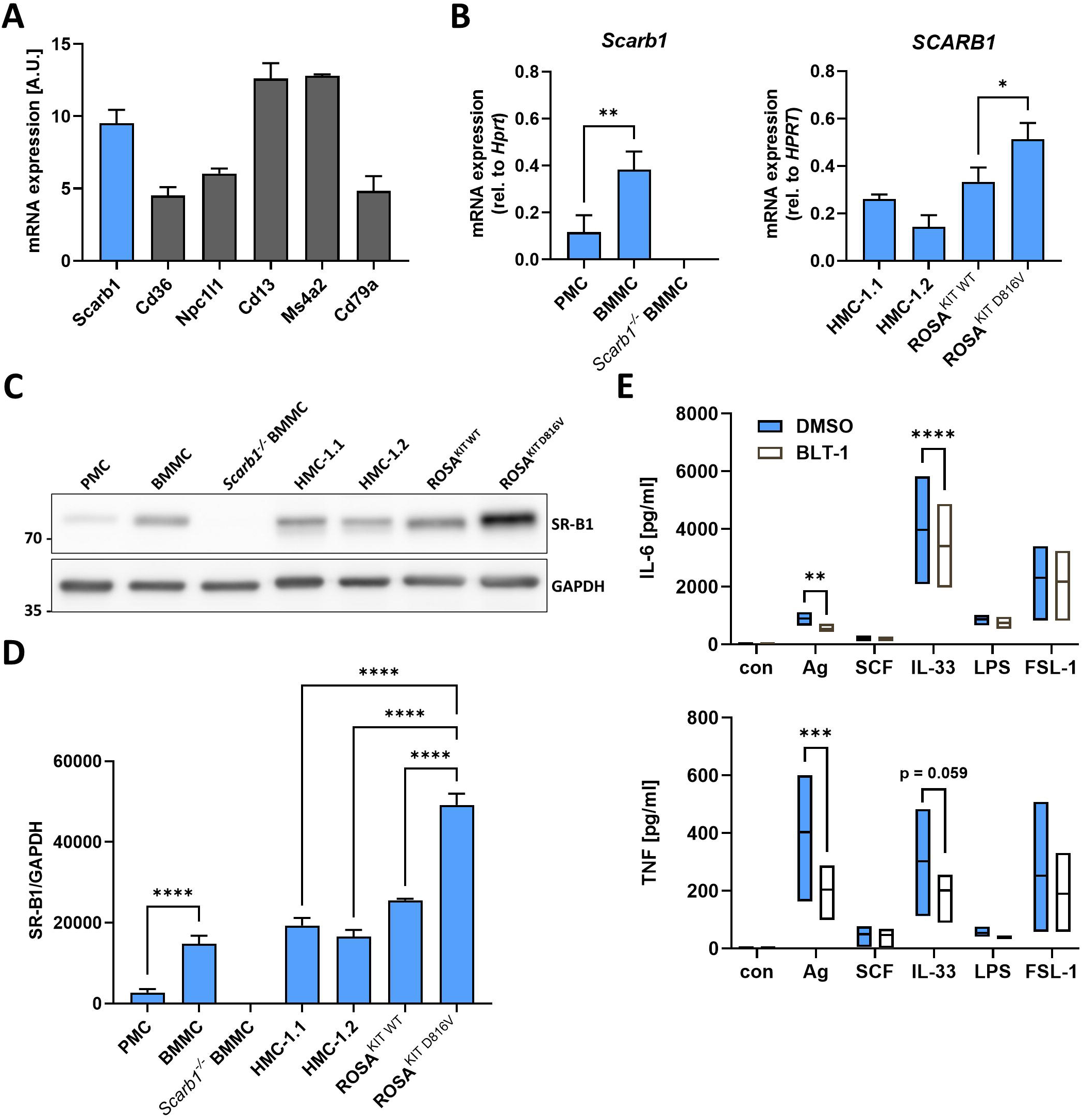
*Scarb1/SCARB1* is expressed in different MC types and inhibition of SR-BI reduces Ag- and IL-33-induced pro-inflammatory cytokine production. **(A)** mRNA expression data of different Ezetimibe targets (*Scarb1*, *Cd*36, *Npc1l1*, *Cd13*) obtained from transcriptomic analysis. Expression of *Ms4a2* and *Cd79a* served as positive and negative controls, respectively. **(B)** Verification of *Scarb1* expression in PMCs and BMMCs by qPCR. *Scarb1^-/-^* BMMCs served as negative control (n=3, left). qPCR analysis of SCARB1 expression in human MC lines HMC-1.1, −1.2, ROSA^KIT WT^ and ROSA^KIT D816V^ (n=3, right). **(C)** Representative Western blot analysis of SR-BI protein expression in murine and human MCs. GAPDH served as loading control. **(D)** Quantification of SR-BI protein expression data by comparing SR-BI expression to GAPDH (n=3). **(E)** WT BMMCs were pre-treated with the SR-BI selective inhibitor BLT-1 [3 µM] for 1 hour. BMMCs were then stimulated with Ag [20 ng/ml], SCF [100 ng/ml], IL-33 [10 ng/ml], LPS [1 µg/ml] or FSL-1 [1 µg/ml] for 3 hours. The released amounts of IL-6 (left) and TNF (right) were determined by ELISA (n=3). Data are shown as mean +SD. Floating bars indicate range between the minimal and maximal values of the independent replicates, the line indicates the mean value. **(B)** and **(D)** were analyzed by ordinary one-way ANOVA followed by Tukey test to correct for multiple comparisons. **(E)** was analyzed by two-way ANOVA followed by Sídák multiple comparisons test. * *p*<0.05, ** *p*<0.01, *** *p*<0.001, **** *p*<0.0001.

### SR-BI is localized on the surface of MCs and co-internalizes together with the FcεRI upon Ag stimulation

To mediate cholesterol transport from HDL to the PM and vice versa, a cell surface localization of SR-BI is necessary. We hypothesized that SR-BI also needs to localize to the cell surface to impact on FcεRI signaling. Therefore, we assessed cell surface localization of SR-BI in different human and murine MCs by flow cytometry. Interestingly, cell surface localization did not fully correlate with total *Scarb1*/SR-BI expression. Although we measured less total *Scarb1* and SR-BI expression in PMCs in relation to BMMCs (Fig. 2B), cell surface localization of SR-BI was similar in both cell types (Fig. 3A & B). This might indicate that additional intracellular pools of SR-BI exist in BMMCs, while in PMCs SR-BI exclusively localizes to the PM. Accordingly, though *SCARB1* expression in HMC-1.1 and −1.2 was similar, cell surface SR-BI localization was higher in HMC-1.2 cells (Fig. 3A & B) suggesting that the different cellular SR-BI pools might by cell type specific. We assumed that SR-BI-dependent regulation of the initial FcεRI activation requires spatial proximity between the two receptors in the PM. Consequently, we measured the co-internalization of SR-BI and FcεRI upon Ag stimulation by staining for PM-localized SR-BI and FcεRI. In WT BMMCs, SR-BI and FcεRI co-internalized with the same time kinetics upon FcεRI crosslinking, corroborating proximity of the receptors (Fig. 3C & D). *Ship1^-/-^* BMMCs have a hyperactive phenotype due to increased PI3K signaling and reportedly degranulate in response to SCF (40). However, SCF-dependent degranulation in *Ship1^-/-^* BMMCs did not lead to co-internalization of SR-BI together with KIT across the observed period. Though the representative FACS histograms give the impression of SR-BI internalization within the first 10 minutes, the quantified MFIs of three independent experiments did not significantly decrease across all time points evaluated (Fig. 3E & F). Thus, we assumed that SR-BI associates and specifically co-internalizes with the FcεRI upon its activation by Ag.

**Fig. 3:**
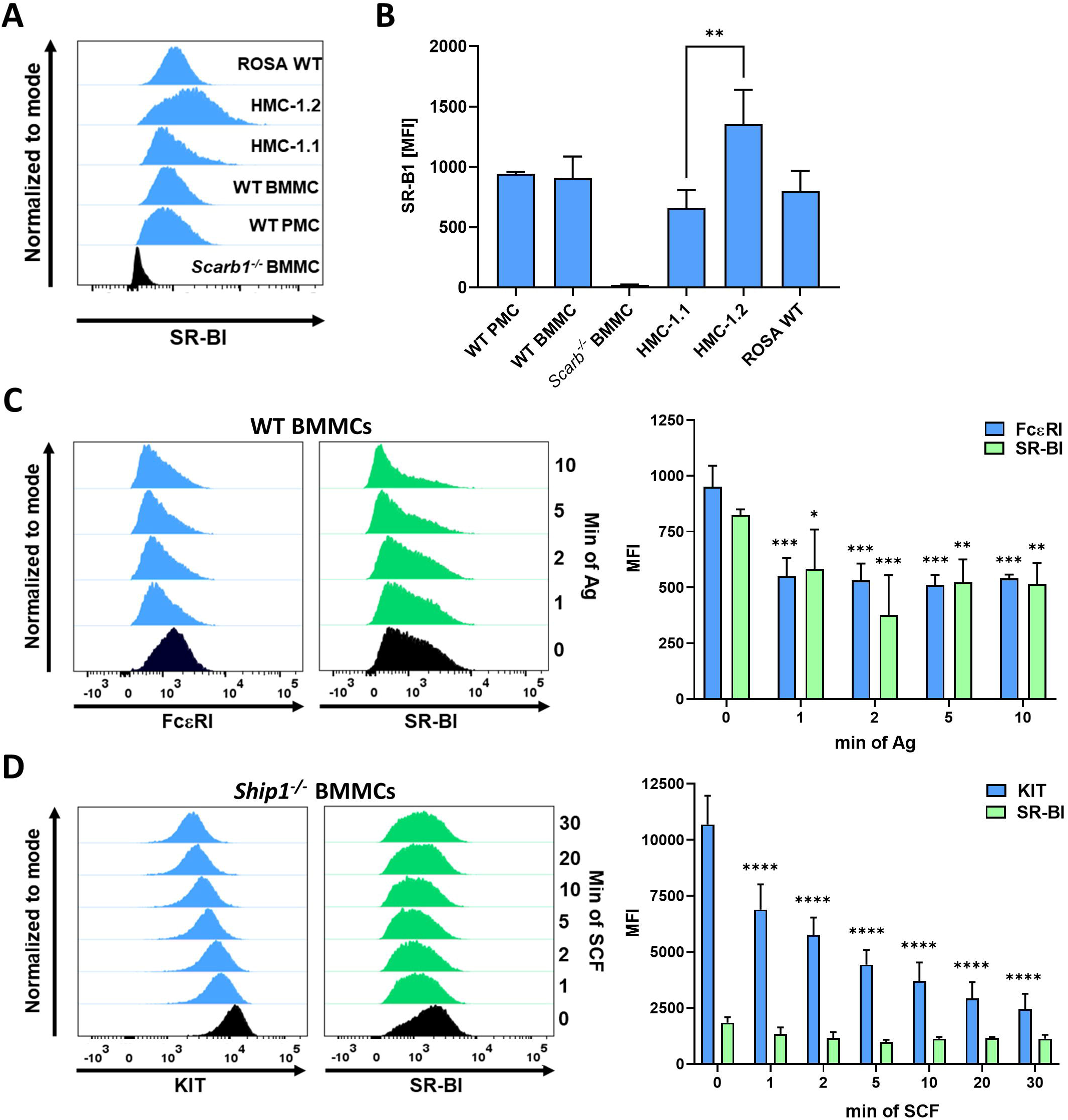
SR-BI localizes to the plasma membrane in MCs and functionally associates with the FcεRI. **(A)** Representative FACS histogram showing cell surface localization of SR-BI in different MC types determined by flow cytometry using an anti-SR-BI antibody specifically recognizing both human and murine SR-BI. **(B)** Quantification of the flow cytometry data shown in **(A)** (n=3). **(C)** WT BMMCs were stimulated with 20 ng/ml Ag for indicated time points. FcεRI and SR-BI expression at the cell surface was determined by flow cytometry. Representative histograms (left) and quantification by MFI (n=3, right) showing receptor internalization. **(D)** *Ship1^-/-^* BMMCs were stimulated with SCF [100 ng/ml] for indicated time points. KIT and SR-BI expression at the cell surface were determined by flow cytometry. Representative histogram (left) and quantification (right) of n=3 experiments. Data are shown as mean +SD. **(B)** was analyzed by ordinary one-way ANOVA followed by Tukey test to correct for multiple comparisons. **(C)** and **(D)** were analyzed by two-way ANOVA followed by Dunnett multiple comparisons test. Asterisks indicate significant differences relative to unstimulated control conditions within the same group. * *p*<0.05, ** *p*<0.01, *** *p*<0.001, **** *p*<0.0001.

### Knockout of *Scarb1* particularly affects synergistically enhanced cytokine production in response to Ag together with SCF, IL-33, PGE_2_ or adenosine

Encouraged by the findings that SR-BI is expressed in MCs, and that the SR-BI inhibitor BLT-1 showed effects on Ag-dependent MC activation, we further examined the mechanistic role of SR-BI on MC activation by analyzing WT BMMCs and BMMCs derived from *Scarb1^-/-^* mice (36). Differentiation of BMMCs from bone-marrow cells was not affected by *Scarb1* deficiency. After 4 weeks of MC differentiation from bone marrow-derived stem cells in the presence of IL-3, cell populations of WT and *Scarb1^-/-^* BMMCs appeared similar in flow cytometry (suppl. Fig. 2A). The amount of cell surface localized FcεRI, KIT, CD13 and ST2 was not affected by *Scarb1* deficiency (suppl. Fig. 2B & C). We analyzed the Ag-triggered degranulation response by determination of β-hexosaminidase release and externalization of the lysosomal marker LAMP1 in WT and *Scarb1^-/-^* BMMCs. Corroborating the findings obtained with BLT-1, SR-BI had no influence on FcεRI-dependent β-hexosaminidase release (suppl. Fig. 3A). Further, quantification of LAMP1 by MFI at the cell surface upon FcεRI stimulation did not reveal differences between WT and *Scarb1*-deficient cells (suppl. Fig. 3B) and BMMCs degranulated to a similar extent and to comparable numbers irrespective of SR-BI presence or absence (suppl. Fig. 3C-E). This suggests that SR-BI is not important for the rapid FcεRI-dependent MC activation processes occurring within seconds to minutes. Therefore, we concentrated on the later pro-inflammatory cytokine response of MCs and determined transcriptional induction of pro-inflammatory mRNAs by qPCR and production/secretion of IL-6 and TNF by ELISA. Accordingly, we stimulated WT and *Scarb1*^-/-^ BMMCs with Ag and – as the effects seen with BLT-1 were rather weak compared to the DMSO control – we additionally pre-stimulated the cells with increasing concentrations of SCF 5 minutes prior to Ag stimulation to analyze synergistic MC activation. Intriguingly, while the detected amounts of *Il6* and *Tnf* mRNA were comparable for both WT and *Scarb1*-deficient BMMCs upon Ag stimulation alone, pre-stimulation with increasing concentrations of SCF led to a stronger increase of pro-inflammatory mRNAs in WT than in *Scarb1^-/-^* BMMCs. This difference became meaningful when the SCF-Ag co-stimulation provoked a significant increase in the levels of *Il6* and *Tnf* mRNAs relative to Ag stimulation alone (Fig. 4A). Similarly, on protein level *Scarb1^-/-^* BMMCs did not exhibit significantly reduced IL-6 and TNF production at the tested Ag concentrations. However, 5 minutes pre-stimulation of BMMCs with a low dose of SCF followed by sub-optimal Ag stimulation led to a synergistically enhanced production of IL-6 and TNF, which was significantly reduced in *Scarb1*-deficient BMMCs (Fig. 4B). When optimal doses of SCF (100 ng/ml) and Ag (20 ng/ml) were used for stimulation to yield a maximal cytokine response, there was a tendency of reduced IL-6 and TNF production for the individual stimuli in *Scarb1*-deficient compared to WT BMMCs and this difference was enhanced to a significant level upon co-stimulation SCF and Ag (suppl. Fig. 4A & B), which would imply that both receptors have a certain SR-BI dependence. Additionally, this attenuated synergistic effect in *Scarb1*-deficient BMMCs was not limited to combined SCF/Ag stimulation. We found similar effects when triggering different classes of receptors that are all reported to act synergistically with the FcεRI in terms of MC effector functions. Thus, *Scarb1*-deficient BMMCs produced less pro-inflammatory cytokines upon co-stimulation of IL-33 (ST2), adenosine (A_3_ receptor) and PGE_2_ (EP3 receptor) with Ag (Fig. 4C-E), while cytokine production upon individual receptor stimulation remained unaffected. In summary, this indicates that SR-BI positively regulates the synergistic enhancement of pro-inflammatory cytokine production in a FcεRI-dependent manner. Of note, the type of co-stimulatory receptor did not have an influence on SR-BI-mediated effects as we observed similar differences between WT and *Scarb1^-/-^* BMMCs for activation of RTKs, cytokine receptors and GPCRs as co-stimulatory signals.

**Fig. 4:**
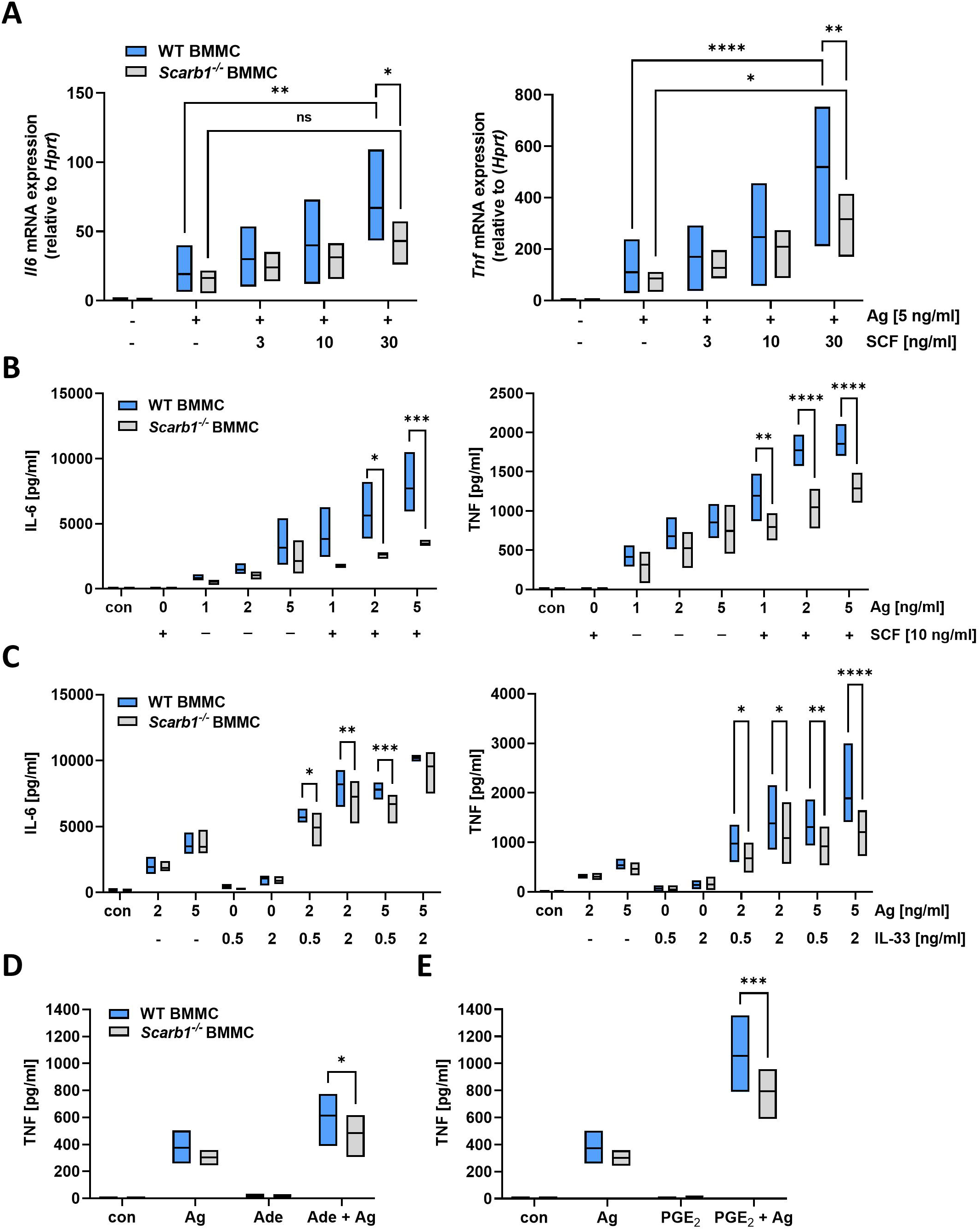
*Scarb1* deficiency particularly suppresses synergistic MC activation. **(A)** WT and *Scarb1^-/-^* BMMCs were left unstimulated, stimulated with 5 ng/ml Ag or stimulated with indicated concentration of SCF for 5 min followed by the addition of 5 ng/ml Ag for 90 minutes. mRNA expression of *Il6* and *Tnf* was analyzed by qPCR. *Hprt* served as housekeeping gene to which expression of other genes was normalized to according to the ΔΔC_T_ method. (n=3) **(B)** WT and *Scarb1^-/-^* BMMCs were left untreated or stimulated with indicated concentrations of either SCF, Ag or the combination of both stimuli with a short SCF pre-stimulation of 5 minutes for a total duration of 3 hours. Secreted IL-6 (left) and TNF (right) was determined from cell supernatants by ELISA (n=3). **(C)** WT and *Scarb1^-/-^* BMMCs were left untreated or stimulated with indicated concentrations of either IL-33, Ag or the combination (5 minutes IL-33 pre-stimulation) for 3 hours. Secreted IL-6 (left) and TNF (right) was determined from cell supernatants by ELISA (n=4). **(D)** WT and *Scarb1^-/-^* BMMCs were left untreated or stimulated with either Ag [5 ng/ml], adenosine [30 µM] or the combination for 3 hours (n=4). Adenosine was added 5 minutes before Ag stimulation. Secreted TNF was determined by ELISA. **(E)** Same experimental setup as in **(D)**, but cells were stimulated with Ag [5 ng/ml, PGE_2_ [1 µM, 5 minutes pre-stimulation] or the combination (n=4). Floating bars indicate range between the minimal and maximal values of the independent replicates, the middle line indicates the mean value. Two-way ANOVA followed by Sídák multiple comparisons test. * *p*<0.05, ** *p*<0.01, *** *p*<0.001, **** *p*<0.0001.

### *Scarb1*-deficient BMMCs have less focally enriched PM cholesterol, which can be mimicked by MβCD and affects PLCγ1 and PKB activation

Based on the broad effects *Scarb1*-deficiency displayed on synergistic cytokine production, we hypothesized that SR-BI-mediated regulation of the MC response might be on the level of receptor activation or immediate downstream. As we did not see effects on one specific type of receptor and knowing that SR-BI facilitates cholesterol shuttling, we assumed that the PM structure and cholesterol-rich nanodomains may influence the synergistic cytokine release. To address this, we stained WT and *Scarb1^-/-^* BMMCs with fluorescently-labelled cholera toxin B (CTB) and the D4 domain of perfringolysin O, which are specific probes for the presence of ganglioside M1 (GM1) and focally enriched cholesterol regions with cholesterol >40 mol% in the PM, respectively (41,42,54). Remarkably, we found that under resting conditions specifically the GFP-D4 staining in *Scarb1^-/-^* BMMCs was reduced compared to the corresponding WT cells. By contrast, the CTB fluorescence was similar between the two genotypes (Fig. 5A). Quantification of the fluorescence intensity in flow cytometry revealed that *Scarb1*-deficient cells had less focally enriched cholesterol in the PM (Fig. 5B). Still, the total cellular cholesterol content (including esterified and unesterified cholesterol) was not affected by the absence of SR-BI but could be drastically reduced in both WT and *Scarb1^-/-^* BMMCs by applying the cholesterol extracting agent MβCD (suppl. Fig. 5A). We verified by FACS analysis that the applied MβCD concentration had no cytotoxic effect as we could not see any shift in the cell population pointing to cell death (suppl. Fig. 5B). Additionally, we corroborated the specificity of the D4 probe for cholesterol by either reducing cholesterol with MβCD or supplementing the cell with MβCD-cholesterol (MβCD-Chol). In flow cytometry we ascertained that the D4 fluorescence decreased by MβCD treatment and increased by adding MβCD-Chol, while there was no change in CTB binding (suppl. Fig. 5C & D). The above-mentioned data suggest that the distribution of cholesterol within the PM might be regulated by SR-BI. Consequently, we asked which pathways downstream of the FcεRI in MCs are dependent on cholesterol and could be disturbed by cholesterol deficiency in cholesterol-rich nanodomains in the PM. Therefore, we pre-treated BMMCs with MβCD for a short time (2 to 5 minutes) to reduce cholesterol in the PM and then stimulated the cells with Ag. Importantly, cholesterol removal strongly reduced the activation of PLCγ1 and the PI3K-PKB pathway, while MAPK pathways were not influenced (Fig. 5C & D). In brief, we could show that the lack of SR-BI leads to reduced cholesterol abundance in cholesterol-rich regions within the PM, which we could mimic using MβCD resulting in reduced activation of PLCγ1 and PKB.

**Fig. 5:**
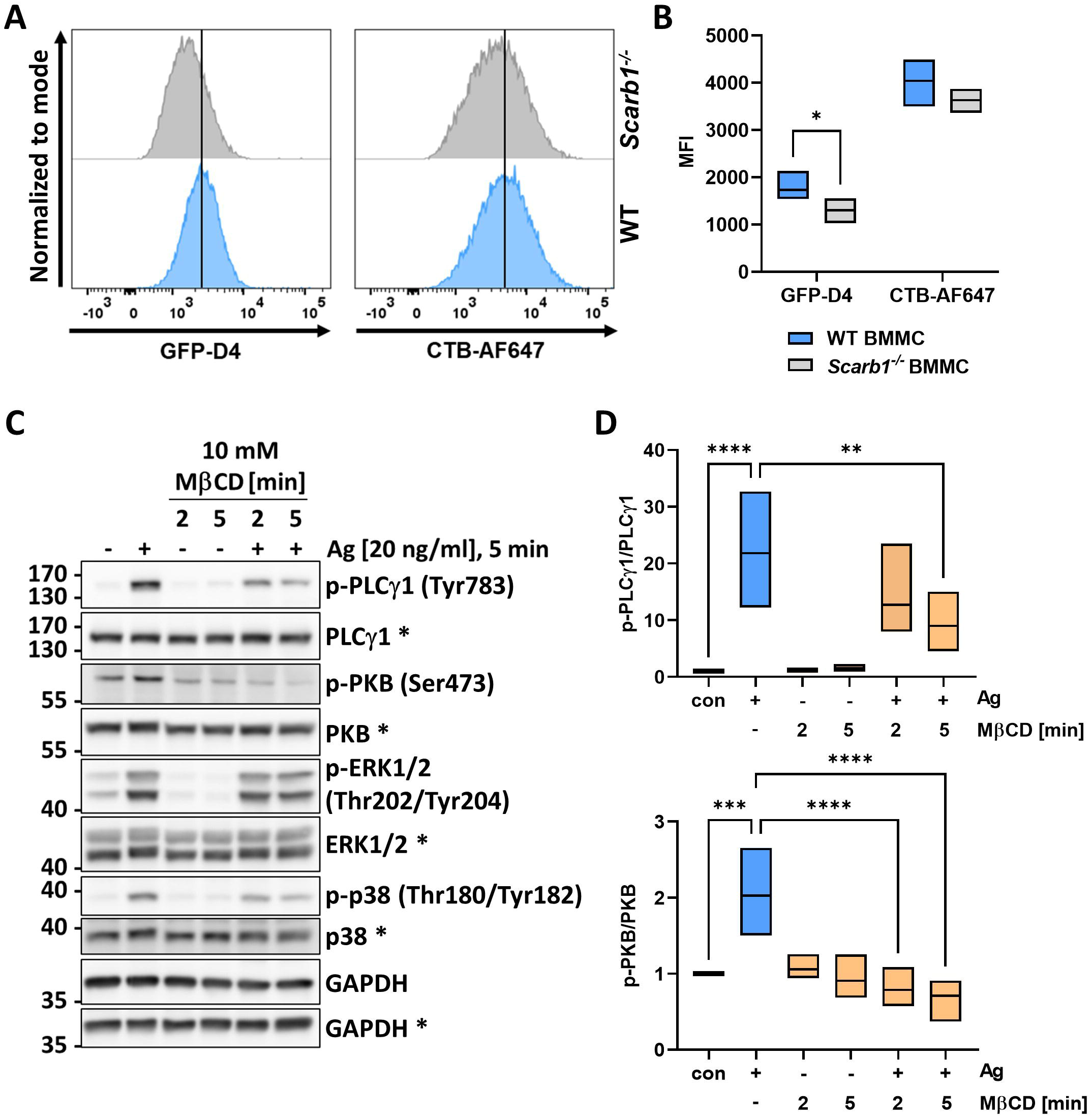
*Scarb1*-deficiency affects mRNA expression data of different mRNA expression data of different membrane impacting on the PI3K-PKB and PLCγ1 pathways. **(A)** Representative histogram of WT and *Scarb1^-/-^* BMMCs stained with the cholesterol- and GM1-specific probes GFP-D4 and CTB coupled to AlexaFluor647, respectively. **(B)** Quantification of n=5 experiments as depicted in **(A)**. **(C)** WT BMMCs were pre-treated with 10 mM MβCD for indicated time and subsequently stimulated or not with 20 ng/ml Ag for 5 minutes. Activation of the PLCγ1, PKB, and MAPK pathways ERK1/2 and p38 was analyzed by phospho-specific antibodies in Western blot. **(D)** Densitometry analysis of p-PLCγ1 and p-PKB in relation to the respective total protein amounts (n=4). Floating bars indicate range between the minimal and maximal values of the independent replicates, the line indicates the mean value. **(B)** was analyzed by multiple unpaired t tests and corrected for multiple comparisons according to the Holm-Sídák method. **(D)** was analyzed by ordinary one-way ANOVA followed by Dunnett test to correct for multiple comparisons. * *p*<0.05, ** *p*<0.01, *** *p*<0.001, **** *p*<0.0001.

### Cholesterol supports the synergistic response to SCF/Ag co-stimulation through PLCγ1 activation

Finally, we focussed on the question how cholesterol or SR-BI can regulate the synergistic MC responses on a signaling level. To investigate this, we focused on the synergism between SCF and Ag as we saw the strongest effects in the previous ELISA measurements when comparing WT and *Scarb1*-deficient BMMCs. In previous experiments, we used MβCD for cholesterol depletion. However, MβCD constitutes a harsh method to disrupt cholesterol-rich regions, which can have several unwanted side effects and may lead to apoptotic cell death if inappropriately applied (55–57). Hence, we used cholesterol oxidase (Chol. ox.) to deplete more specifically cholesterol from lipid rafts by Chol. ox.-catalyzed oxidization of free cholesterol to cholestenone (58). As shown in Fig. 6A, we pre-treated WT BMMCs with H_2_O or Chol. ox. and then stimulated with SCF, Ag, or the combination (5 minutes SCF pre-stimulation before addition of Ag) of both stimuli. Analysis of activated signaling pathways revealed that activation of PLCγ1 was enhanced when KIT and FcεRI were co-activated in relation to SCF or Ag applied alone. Chol. ox. pre-treatment specifically reduced phosphorylation of PLCγ1 at Tyr783 under co-stimulatory conditions, while PLCγ1 activation upon SCF and Ag stimulation alone was comparable between H_2_O and Chol. ox.-treated WT cells. Additionally, the attenuated activation of the PI3K-PKB pathway seen after MβCD treatment was not present after Chol. ox. addition. Notably, PKB phosphorylation was not enhanced by co-stimulation with SCF and Ag, but rather slightly reduced compared to SCF-induced PKB activation. Finally, activation of MAPK pathways was not affected by cholesterol oxidation suggesting no dependence on the presence of cholesterol in the PM (Fig. 6A & B). To validate these findings, we repeated this experiment and used Ezetimibe, which in initial experiments showed strong inhibitory effects on MC effector functions. We hypothesized and in the beginning validated that Ezetimibe, like MβCD, would also show side effects, yet, as a licensed cholesterol absorption inhibitor, it should phenocopy some of the results obtained with MβCD and Chol. ox.. Indeed, pre-treatment of WT BMMCs with Ezetimibe strongly inhibited the activation of PLCγ1 in response to SCF/Ag co-stimulation, while PLCγ1 phosphorylation was not affected upon individual KIT or FcεRI activation (suppl. Fig. 6A & B). Moreover, Ezetimibe also reduced activation of the MAPK p38, but this effect only occurred when BMMCs were stimulated with either SCF or Ag alone. As already described in Fig. 1D, we again noticed an Ezetimibe-mediated reduced activation of the PI3K-PKB pathway in Ag-stimulated cells using phosphorylation of Ser473 of PKB as readout. However, Ezetimibe did not impact on SCF-induced and only slightly on the synergistic SCF/Ag-induced PKB activation (suppl. Fig. 6A & B). This indicates that Ezetimibe as cholesterol-lowering drug in a similar way as MβCD and Chol. ox. treatment attenuates synergistic MC responses as a result of KIT and FcεRI co-activation specifically by reducing PLCγ1 and/or PKB phosphorylation. To ultimately not only emphasize on the role of cholesterol in synergistic MC activation, but also to implicate SR-BI in this mechanism we aimed at verifying whether SR-BI inhibition would have similar consequences as Chol. ox.-mediated cholesterol depletion. Consequently, we used WT BMMCs and pre-treated with either DMSO as control or BLT-1 to inhibit cholesterol shuttling via SR-BI and subsequently stimulated with (sub-)optimal concentrations of Ag alone or in combination with a SCF pre-stimulation for 5 minutes prior to Ag addition. As before, we analyzed signaling pathway activation by Western blot. Intriguingly, similar to what we observed upon cholesterol depletion using Chol. ox., phosphorylation of PLCγ1 at Tyr783 was drastically reduced in BLT-1-treated BMMCs after co-stimulation with SCF and Ag (20 ng/ml) (Fig. 6C & D). Interestingly, a lower Ag concentration (2 ng/ml) was not enough to boost PLCγ1 phosphorylation in DMSO-treated cells, which we only observed after stimulation with 20 ng/ml suggesting that only the latter concentration elicits a proper synergistic response (Fig. 6C & D). Akin to the signaling analysis upon Chol. ox. treatment, activation of PKB measured by analyzing phosphorylation at Ser473 was not influenced by BLT-1 treatment and, moreover, this phosphorylation was not increased by combined activation of KIT and FcεRI in relation to stimulation with SCF or Ag alone (Fig. 6C &D). Again, activation of MAPK pathways p38 and ERK1/2 was not affected by BLT-1 treatment. Hence, as we see analogous effects specifically upon KIT- and FcεRI-dependent co-activation of WT BMMCs when we pre-treat the cells with cholesterol lowering agents (Ezetimibe, MβCD, Chol. ox) or inhibit SR-BI by BLT-1, we propose that SR-BI-mediated regulation of PM cholesterol homeostasis is crucial for synergistic MC activation. Of note, disruption of this regulatory process might disturb the synergism between KIT and FcεRI signaling, which – based on our findings – is primarily attributed to an enhanced phosphorylation of PLCγ1 that is significantly inhibited by cholesterol depletion and/or SR-BI inhibition.

**Fig. 6:**
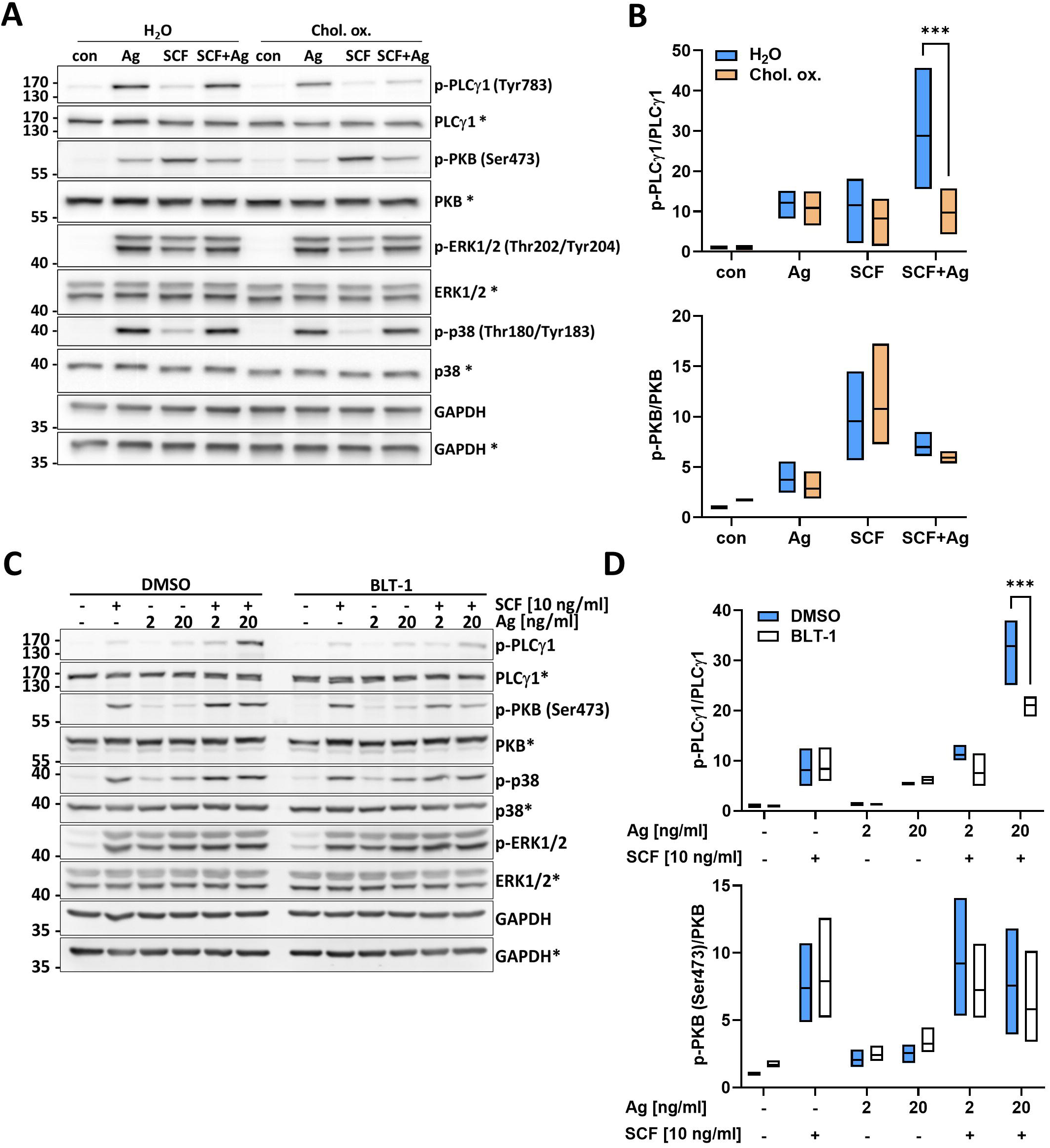
Cholesterol supports the synergistic response to SCF/Ag co-stimulation through PLCγ1 activation. **(A)** Representative Western blot of WT BMMC pre-treated with either H_2_O or Chol. ox. (3 U/ml, 20 min) and subsequently left unstimulated or stimulated with Ag (20 ng/ml, 5 min), SCF (10 ng/ml, 10 min) or the combination of both stimuli (SCF 5 min prior to Ag for 5 min). Activation of signaling pathways was analyzed by Western blot using phospho-specific antibodies recognizing phosphorylation of PLCγ1 (Tyr783), PKB (Ser473), ERK1/2 (Thr202/Tyr204) and p38 (Thr180/Tyr183). The respective amounts of detected total protein served as loading control. Total proteins and phosphorylated proteins were detected on separate membranes, which is indicated by an asterisk next to the protein name. GAPDH served as control to prove equal loading of both membranes. **(B)** Quantification of the signal intensities of phosphorylation of PLCγ1 and PKB in Western blot results from **(A)**. Quantification was performed by densitometry analysis from n=3 independent experiments. **(C)** Representative Western blot of WT BMMCs pre-treated with DMSO or BLT-1 (3 µM) for one hour and subsequently left unstimulated or stimulated with indicated concentrations of Ag for 5 min, 10 ng/ml SCF for 10 min or the combination (5 min SCF pre-stimulation followed by Ag stimulation for 5 min). Activation of signaling pathways was analyzed as in **(A)**. **(D)** Quantification of phosphorylation intensities of PLCγ1 and PKB relative to total protein in Western blots from **(C)**. Densitometry analysis was performed from n=3 independent experiments. Floating bars indicate range between the minimal and maximal values of the independent replicates, the middle line indicates the mean value. Two-way ANOVA followed by Sídák multiple comparisons test. *** *p*<0.001.

### Synergistic activation of human ROSA^KIT WT^ mast cells inhibited by BLT-1

To complete our studies about the involvement of SR-BI and cholesterol in synergistic MC activation, we employed an alternative cellular model to substantiate our findings. As we exclusively used BMMC for our previous experiment, we now switched to the established human MC line ROSA^KIT WT^. As ROSA^KIT WT^ cells are routinely cultivated in SCF-containing medium, we suspected that they are not a suitable model to study the synergism between KIT and FcεRI due to persistent KIT activation under regular culture conditions. However, as we noticed a comparable importance of SR-BI for the other synergistic responses in BMMCs, as well, we focussed on the reaction of ROSA^KIT^ ^WT^ cells to Ag, PGE_2_ or the combination of both stimuli with respect to induction of mRNA expression of pro-inflammatory genes. To address the relevance of SR-BI in ROSA^KIT^ ^WT^ cells for the PGE_2_/Ag synergism, we pre-treated the cells with DMSO as control or BLT-1 for 1 hour prior to stimulation with either PGE_2_ or Ag alone or in combination. As a readout, we determined induction of *IL8*, *TNF* and *CCL1* mRNA expression. Application of PGE_2_ and Ag alone induced *IL8* expression to a similar extent in DMSO- and BLT-1-pre-treated ROSA^KIT WT^ cells. Importantly, we could measure a prominent increase in *IL8* expression in DMSO-pre-treated cells when both stimuli were combined and this enhancement was drastically reduced when SR-BI was inhibited (Fig. 7A). In a similar way, induction of *TNF* expression was specifically attenuated when ROSA^KIT WT^ cells were treated with BLT-1 and co-stimulated with PGE_2_ and Ag, while BLT-1 did not affect the response to each individual stimulus (Fig. 7B). In contrast to *IL8* and *TNF*, induction of *CCL1* expression was not at all affected by SR-BI inhibition, neither by single stimulation nor upon combined application of PGE_2_ and Ag (Fig. 7C). This points towards the hypothesis that enhanced expression of pro-inflammatory genes upon co-stimulation is not overall controlled by SR-BI but that it may depend on the cholesterol sensitivity of pathways necessary for transcriptional induction of respective genes whether or not SR-BI affects their expression.

**Fig. 7:**
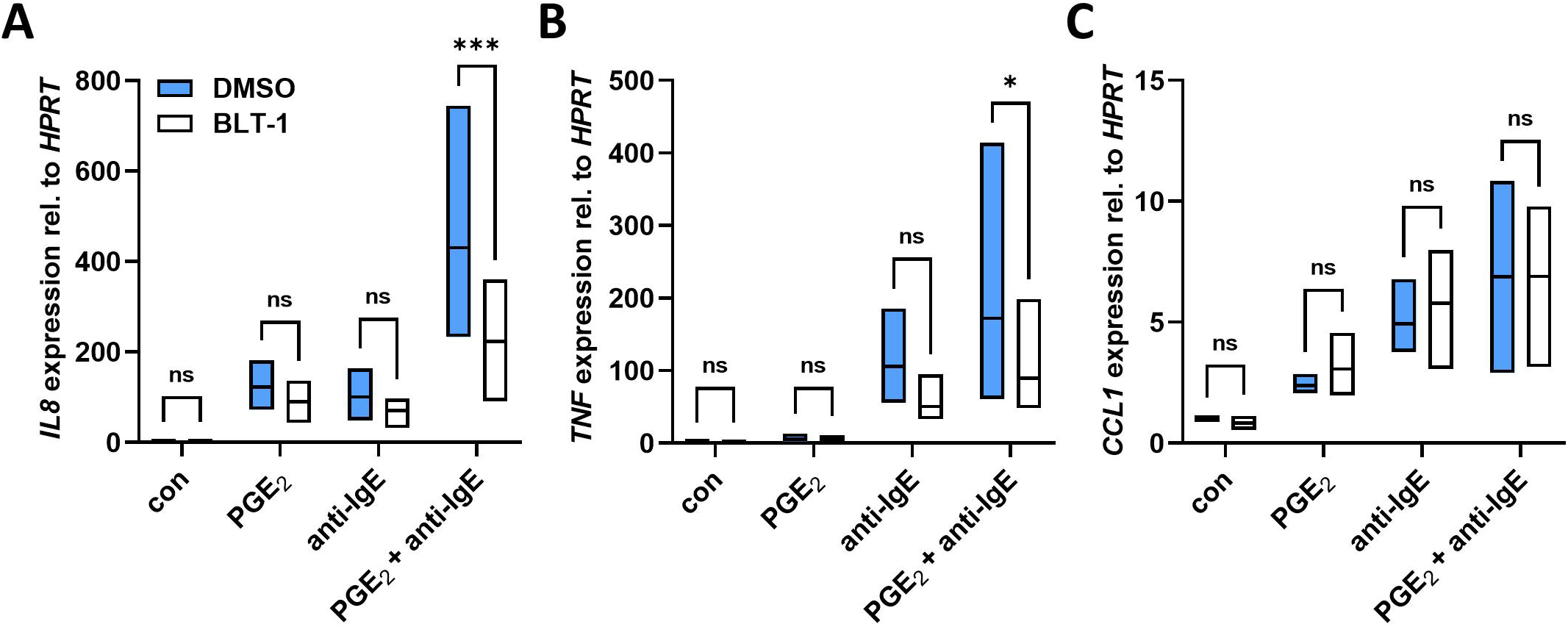
The synergistic response of ROSA^KIT^ ^WT^ cells to PGE_2_/Ag co-stimulation can be inhibited by SR-BI inhibition. **(A)-(C)** ROSA^KIT WT^ cells were cultivated for 4 days in the presence of IL-4 (20 ng/ml) and IgE (1 µg/ml) to upregulate FcεRI expression at the cell surface. Before stimulation, ROSA^KIT WT^ cells were pre-treated with DMSO or BLT-1 (3 µM) for one hour and subsequently left unstimulated or stimulated with PGE_2_ (0.1 µM) for 65 min, anti-IgE (0.5 µg/ml) for 60 min or the combination (5 min PGE_2_ pre-stimulation followed by 60 min anti-IgE stimulation). mRNA expression of IL8 **(A)**, TNF **(B)** and CCL1 **(C)** was determined by qPCR analysis using HPRT as housekeeping gene to which expression of genes of interests was normalized to by the ΔΔC_T_ method (n=4). Floating bars indicate range between the minimal and maximal values of the independent replicates, the middle line indicates the mean value. Two-way ANOVA followed by Sídák multiple comparisons test. Ns *p*>0.05, * *p*<0.05, *** *p*<0.001.

## Discussion

For the common IgE-dependent type I hypersensitivity reactions, MCs are the central effector cells initiating allergic inflammation by the release of numerous pro-inflammatory molecules and proteins (1). The secretion depends on the activation of IgE-bound FcεRI through crosslinking of multiple receptors by multivalent Ag (2). Co-receptors activated by respective ligands present in the surrounding microenvironment can synergize with FcεRI crosslinking to boost MC activation in a para- or autocrine fashion. Here, we describe a role for SR-BI in regulating the raft-associated cholesterol content necessary for proper enhancement of pro-inflammatory cytokine production. We could reveal that SR-BI is widely expressed in different murine and human MC types independent of the presence of FcεRI expression and functionality, which is absent in the leukemic MC lines HMC-1.1 and −1.2 (59). Notably, SR-BI expression was even increased in HMC-1.2 and ROSA^KIT D816V^ cells arguing for a possible MC leukemia-promoting role.

We continued to concentrate on the role of the PM cholesterol homeostasis regulated by SR-BI impacting on FcεRI-dependent signaling processes in MCs. SR-BI is known for its ability to mediate cellular HDL-derived cholesterol uptake and also release onto extracellular acceptors (22,27,28). This implicates that SR-BI functions in regulating the cellular cholesterol abundance. Importantly, SR-BI possesses a cholesterol-sensing domain contributing to monitor the actual cholesterol concentration and respond to changes of the cholesterol content in the PM (26,60). We could show that SR-BI deficiency resulted in a significantly reduced amount of cholesterol specifically in PM cholesterol-rich regions shown by reduced GFP-D4 binding to the surface of *Scarb1^-/-^* BMMCs in relation to WT BMMCs. Intriguingly, a study by Kellner-Geibel et al. (27) revealed that expression of SR-BI in COS-7 cell led to a re-distribution of the cellular cholesterol pool. In this study, SR-BI was especially implicated in the enrichment of cholesterol in caveolae regions within PM that serve as membrane domains enabling influx and efflux of cholesterol between cells and extracellular acceptors. Similar to lipid rafts, also caveolae are rich in cholesterol, which is thought to contribute to their structural integrity (61,62). Hence, it is likely that SR-BI in a comparable way might also regulate the abundance of cholesterol in lipid raft regions and thereby impact on the spatial organization of signaling processes. Of note, the presence of GM1 was not affected by SR-BI-deficiency suggesting that SR-BI has no influence on other lipid raft constituents. It is also worth mentioning that we could detect less PM cholesterol by GFP-D4 staining in *Scarb1*-deficient cells though the cellular cholesterol homeostasis is tightly regulated by cholesterol-sensing mechanisms within the ER that respond to cholesterol deficiency by upregulating the endogenous cholesterol biosynthesis (63). Hence, we assumed that *Scarb1* knockout BMMCs would have time to adapt and counterbalance the decreased cholesterol abundance in lipid raft regions of the PM, which did, however, not happen. One could speculate that this might have been due to an increase in free cholesterol in non-raft regions because of the missing shuttling of cholesterol into lipid rafts facilitated by SR-BI, which would not result in a general deficiency but a re-distribution of cholesterol in the PM.

Several studies have shown the intricate relationship between cholesterol and the size and integrity of lipid raft domains (64,65). In this regard it has been uncovered that raft-associated cholesterol is required for the localized clustering of PI-4,5-P_2_ in raft regions (66,67). Crucially, PI-4,5-P_2_ is the substrate of both PLCγ1 and PI3K that either hydrolyze PI-4,5-P_2_ to IP_3_ and DAG or phosphorylate PI-4,5-P_2_ to PI-3,4,5-P_3_, respectively. Besides, it was shown only recently that PLCγ1 specifically interacts with PI-4,5-P_2_ via its N-terminal PH and EF hand domains as well as with cholesterol via its C2 and TIM barrel domains, which is important for membrane targeting and catalytic activity (68). This would fit to our results demonstrating that phosphorylation of PLCγ1 and PKB are substantially reduced upon Ag-dependent FcεRI activation when BMMCs were pre-treated with MβCD to extract membrane cholesterol. Apart from its cholesterol-extracting properties, MβCD is a strong lipid raft disrupting agent (69), which would impair membrane targeting of both PKB and PLCγ1 signaling complex formation at the PM (70). Intriguingly, when we applied Chol. ox. instead of MβCD, which specifically depletes cholesterol from raft regions but does not lead to disruption of entire lipid raft structures (35), we observed a decreased PLCγ1 phosphorylation while Ser473 phosphorylation of PKB remained unaffected. Notably, this cholesterol-dependent activation of PLCγ1 did only occur upon synergistic MC stimulation with SCF and Ag as cholesterol oxidation did not impact PLCγ1 activation following individual KIT or FcεRI stimulation in BMMCs. PLCγ1 binds to phosphorylated LAT following FcεRI activation and directly binds to auto-phosphorylated KIT, which both contributes to its membrane recruitment and activation (71,72). Besides, it has been convincingly demonstrated that PKB activation is dependent on the presence of lipid raft structures (73). However, it was presently unknown that enhanced PLCγ1 activation in co-stimulated BMMCs depends on the cholesterol present in lipid raft regions. This underlines the importance of the PM structure for MC signaling initiation and suggests that cholesterol is a critical positive regulatory factor controlling membrane recruitment of signaling proteins. Therefore, it would be tempting to speculate that, under SCF/Ag co-stimulation conditions, PM cholesterol serves as an additional recruitment mechanism for PLCγ1 – apart from binding to phosphorylated LAT and KIT – due to the affinity of the C2 and TIM barrel domains for PM cholesterol. We only measured PLCγ1 phosphorylation as a readout for its catalytic activity but not the recruitment to the PM, which could be disturbed when less cholesterol is present in lipid raft structures. Besides, enhanced phosphorylation of PLCγ1 at Tyr783, which is a readout for its catalytic activity (74), would also suggest an increased Ca^2+^ response. This should accordingly be decreased in *Scarb1^-/-^* BMMCs or in either BLT-1-pre-treated or cholesterol-depleted WT BMMCs, which was not tested in this study.

In BMMCs, the absence of SR-BI significantly diminished IL-6 and TNF production upon Ag co-stimulation with SCF, IL-33, adenosine and PGE_2_. However, pre-treatment of WT BMMCs with a SR-BI selective inhibitor also diminished pro-inflammatory cytokine production in response to FcεRI activation alone. This was dependent on pleiotropic effects of BLT-1 on different signaling pathways (PLCγ1, PKB, p38). Still, this effect was only seen at late time points following Ag stimulation. When we analyzed the effect of BLT-1 also on SCF/Ag co-stimulation, we could phenocopy the reduced PLCγ1 phosphorylation previously noticed with Chol. ox. pre-treatment at earlier time points. SCF, IL-33 as well as adenosine and PGE_2_ employ receptors that function in different ways and activate distinct signaling pathways. Thus, we hypothesized and validated that SR-BI exerts its regulatory effect on synergistic MC activation by controlling the PM-associated cholesterol distribution, which was reduced in PM raft regions of *Scarb1*-deficient BMMCs. Still, the question that arises is why cholesterol is specifically important for the synergistic pro-inflammatory cytokine production of MCs and how PLCγ1 contributes to this synergism. Interestingly, previous studies investigating the signaling mechanisms behind synergistic MC activation described that the synergy between FcεRI and SCF, PGE_2_ or adenosine are either dependent on increased PI3K-PKB or PLCγ1 signaling. Consequently, combined activation of different PI3K isoforms was reported to increase MC activation in response to combined adenosine and Ag stimulation (20) while activation of different PLC isoforms is thought to be responsible for integrating signals stemming from combined PGE_2_ and Ag stimulation (21). Importantly, as activation of both PKB and PLCγ1 are cholesterol-sensitive as shown herein, the previous studies and observations support our view on synergistic MC activation, which implicates cholesterol as another regulatory factor specifically in regulating the integration of signals from co-stimulated receptors. Nevertheless, our data similar to those of Kuehn et al. (21) support the view that PLCs are central to synergistically enhanced pro-inflammatory MC activation at least for the synergy between FcεRI and KIT. As there exists no functional link between GPCR signaling and PLCγ activation, this appears contr-intuitive. However, a study by Patterson et al. (75) found that specifically the SH3 domain of PLCγ is required for a proper Ca^2+^ influx following serotonin and muscarinergic receptor stimulation, which was independent of its catalytic activity. This could provide an explanation how PLCγ1 could influence MC activation in response to co-stimulation of FcεRI and GPCRs. Contrary to PLCs as critical signaling enzymes, one of the earliest studies by Vosseller et al. (16) identified that the Y719 in KIT, which constitutes the binding site of the p85 subunit of PI3K is critically involved in synergistic enhancement of MC effector functions in response to KIT and FcεRI co-stimulation and hence puts combined PI3K activation at the centre of synergism control. However, other reports mainly dealing with synergistic responses in human MCs again confirmed that primarily PLCγ1 activation and the ensuing increased Ca^2+^ response are fundamental to the FcεRI-based MC synergism (76,77). Of note, using different approaches (Chol. ox. and Ezetimibe pre-treatment as well as BLT-1-mediated SR-BI inhibition) we could provide multi-layered evidence that by impacting on the PM- and raft-associated cholesterol pool we were able to explicitly influence the PLCγ1 activation upon SCF/Ag co-stimulation in BMMCs, which supposedly is responsible for the decreased pro-inflammatory cytokine production in *Scarb1*-deficient MCs.

In studies by Tkaczyk et al. and Hundley et al. (76,77) the authors also focussed on the degranulation response, which in our experiments appeared not to be influenced by SR-BI deficiency or inhibition though it is known that proper FcεRI crosslinking and activation is lipid raft dependent (31,78). Moreover, activation of RTKs, GPCRs and cytokine receptors is thought to be regulated by functional raft domains in the PM (79–81). However, we could only measure a slightly reduced raft-associated cholesterol content in *Scarb1*-deficient BMMCs compared to WT counterparts by GFP-D4 staining, while MβCD treatment, which robustly inhibited PKB and PLCγ1 activation upon Ag stimulation alone drastically diminished the PM cholesterol pool. This could indicate that subtle PM cholesterol changes do not severely affect the response upon individual receptor stimulation but that simultaneous activation of different receptors that all require cholesterol-rich raft domains might amplify the effects of a reduced PM cholesterol pool.

Finally, using human ROSA^KIT WT^ cells, we could in a complementary MC model show, that SR-BI inhibition similarly affects the synergistic cytokine response upon PGE_2_/Ag co-stimulation. *IL8* and *TNF* production could be significantly decreased by BLT-1 pre-treatment upon co-stimulation while there was no effect on *CCL1* production. Interestingly, *IL8* and *TNF* production have been shown to be dependent on NFAT activation, which is activated in a Ca^2+^-dependent way implying a certain PLCγ1 dependence (82). By contrast *CCL1* expression is primarily dependent on MAPK kinase pathway activation (83,84). As we could show that MAPK activation was unaffected by cholesterol depletion and SR-BI inhibition in BMMCs, this provides an explanation why *CCL1* expression in ROSA^KIT WT^ cells was not influenced by BLT-1 pre-treatment. In sum, we herein described that the synergistic pro-inflammatory MC response is dependent on raft-associated PM cholesterol abundance, which is regulated by SR-BI. This novel regulatory mechanism could open up new possibilities for pharmacological intervention to treat unwanted MC activation in allergic patients.

## Data availability statement

The raw data supporting the conclusions of this article will be made available by the authors.

## Author contributions

SC and MK performed experiments, SC analyzed the data. SC and MH conceptualized the study. SC wrote the draft of the manuscript. SC, MA and MH contributed to review and editing. MH contributed to funding acquisition. MA provided ROSA^KIT WT^ and ROSA^KIT D816V^ cells.

## Acknowledgements

We would like to thank Monty Krieger (Massachusetts Institute of Technology) for providing *Scarb1^-/-^* mice as well as Toshihide Kobayashi (University of Strasbourg) and Mitsuhiro Abe (RIKEN Institute) for providing pET28/His6-EGFP-D4 plasmid DNA.

---

**Supplementary fig. 1:**
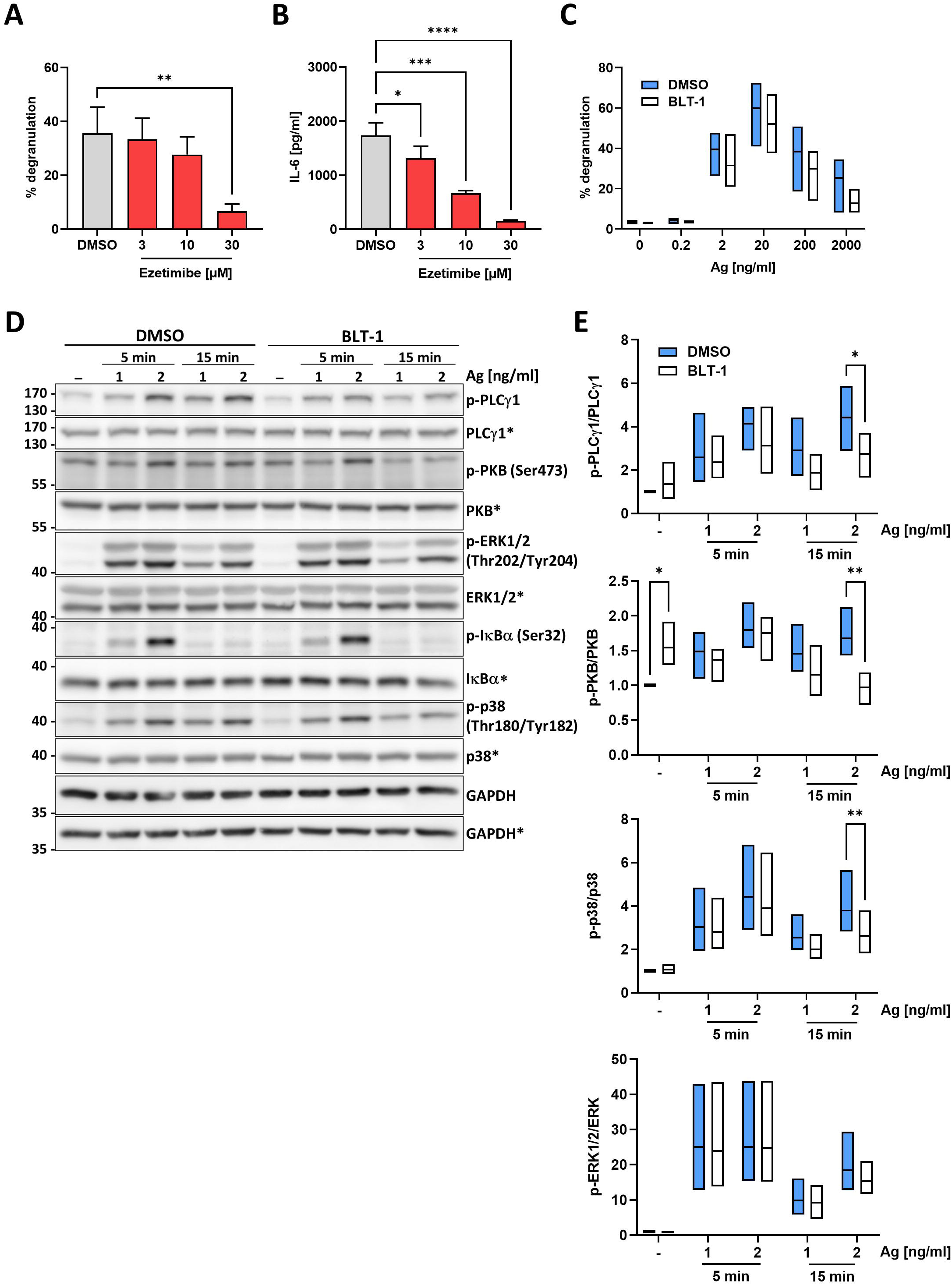
*Scarb1*-deficient cells are sensitive to Ezetimibe treatment and SR-BI inhibition reduces PLCγ1, PKB and p38 activation. **(A)** *Scarb1^-/-^* BMMCs were pre-treated with indicates Ezetimibe concentrations for 1 hour and subsequently stimulated with 20 ng/ml Ag for 30 minutes. Degranulation was determined by β-hexosaminidase assay (n=3). **(B)** Same experimental setup as in **(A)**, but cells were stimulated for 3 hours and secreted IL-6 was measured in cell culture supernatants by ELISA (n=3). **(C)** WT and *Scarb1^-/-^* BMMCs were stimulated with indicated Ag concentrations for 30 minutes and degranulation was determined by b-hexosaminidase assay (n=3). **(D)** Representative Western blot analysis of 1 hour DMSO or BLT-1 [3 µM] pre-treated and Ag stimulated WT BMMCs. Activation of PLCγ1, PKB, MAPK pathways ERK1/2 and p38 and NF-κB was determined. **(E)-(G)** Densitometry analysis of p-PLCγ1, p-PKB and p-p38 in relation to the respective total protein amounts of the Western blot depicted in **(D)** (n=3). Data are expressed as mean +SD or as floating bars indicating minimal, maximal and mean values. **(A)** and **(B)** were analyzed by ordinary one-way ANOVA followed by Dunnett test to correct for multiple comparisons. **(E)-(G)** were analyzed by two-way ANOVA followed by Sídák multiple comparisons test. * *p*<0.05, ** *p*<0.01, *** *p*<0.001, **** *p*<0.0001.

**Supplementary figure 2:**
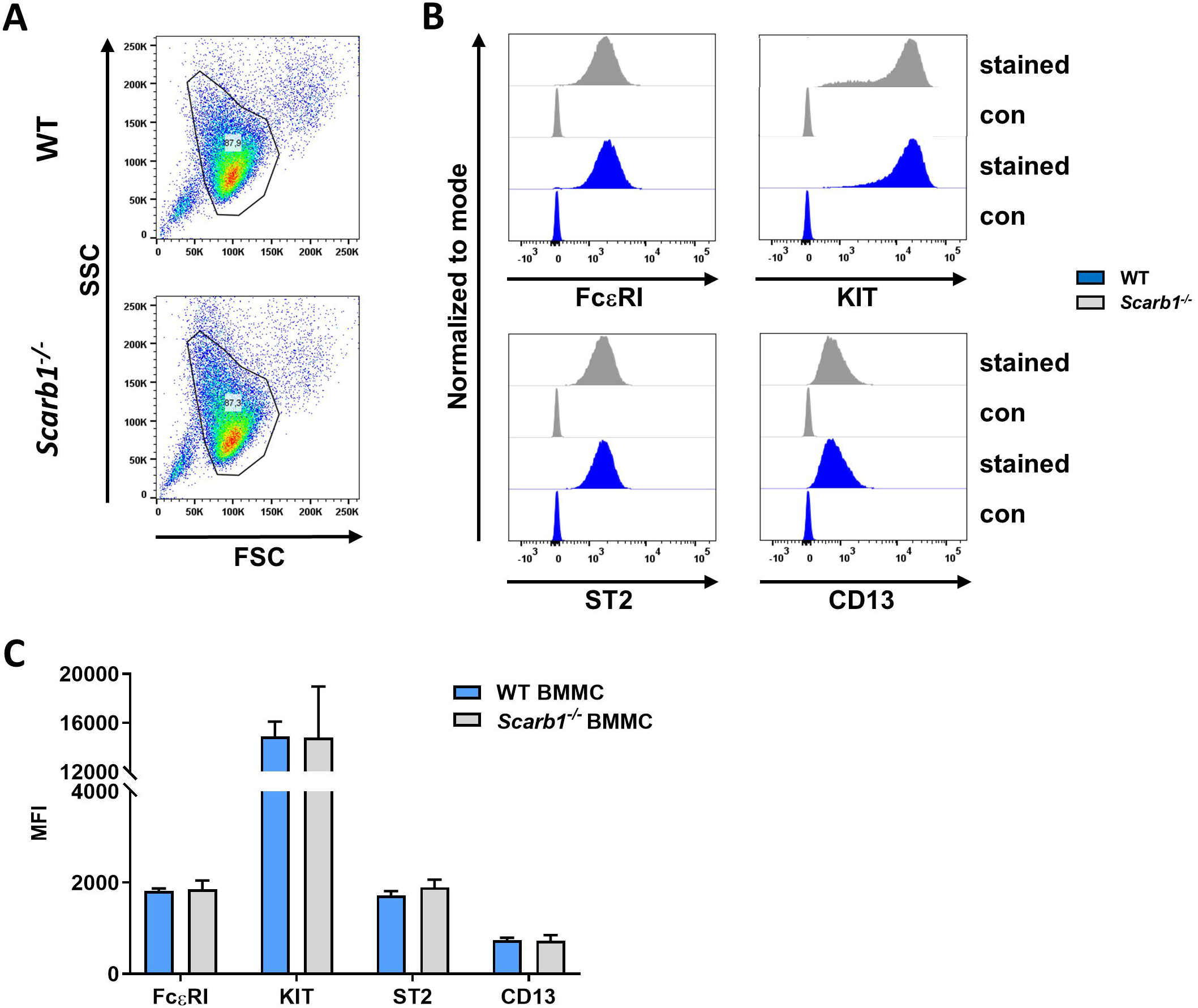
*Scarb1*-deficiency did not affect BMMC differentiation. **(A)** Representative FACS dot plots showing BMMC populations of WT and *Scarb1^-/-^* BMMCs differentiated for 4 weeks in the presence of IL-3. **(B)** Representative histograms of MC surface marker expression (FcεRI, KIT, ST2, CD13) in WT and *Scarb1^-/-^* BMMCs after 4 weeks of differentiation. **(C)** Quantification of data depicted in **(B)** (n=4). Data are expressed as mean +SD.

**Supplementary figure 3:**
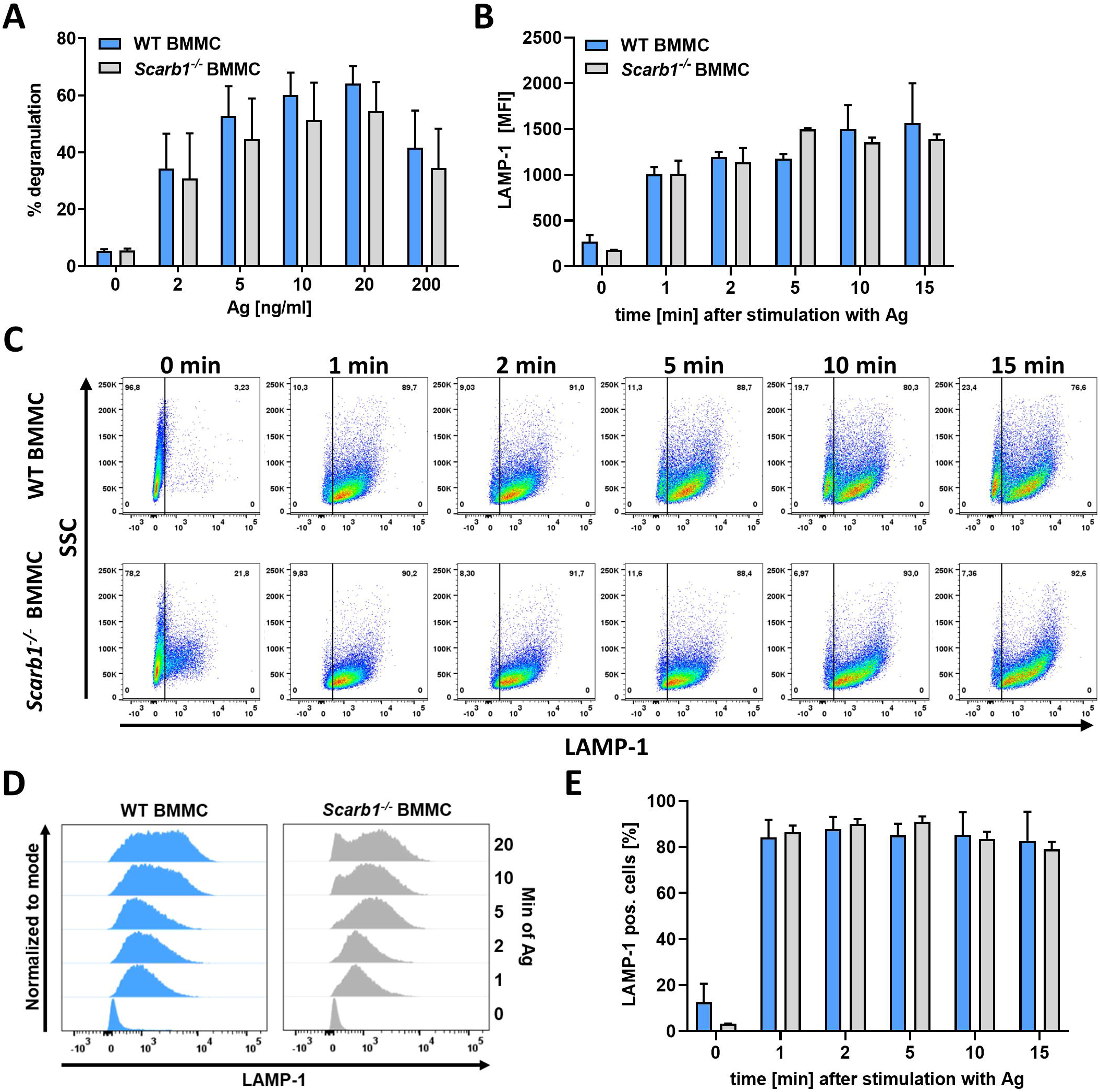
Mast cell degranulation is not regulated by SR-BI. **(A)** WT and *Scarb1^-/-^* BMMCs were stimulated with indicated concentrations of Ag for 30 minutes and degranulation was determined by β-hexosaminidase assay (n=5). **(B)** WT and *Scarb1^-/-^* BMMCs were stimulated with 20 ng/ml Ag for indicated time and externalization of the granule marker LAMP-1 was determined by flow cytometry analysis quantified as MFI (n=3). **(C)** Representative dot plots of LAMP1 assay performed with WT and *Scarb1^-/-^* BMMCs. **(D)** Representative histograms of LAMP1 assay performed with WT and *Scarb1^-/-^* BMMCs. **(E)** Quantification of LAMP1 positive cells in % after different stimulation times with Ag [20 ng/ml] (n=3).

**Supplementary figure 4:**
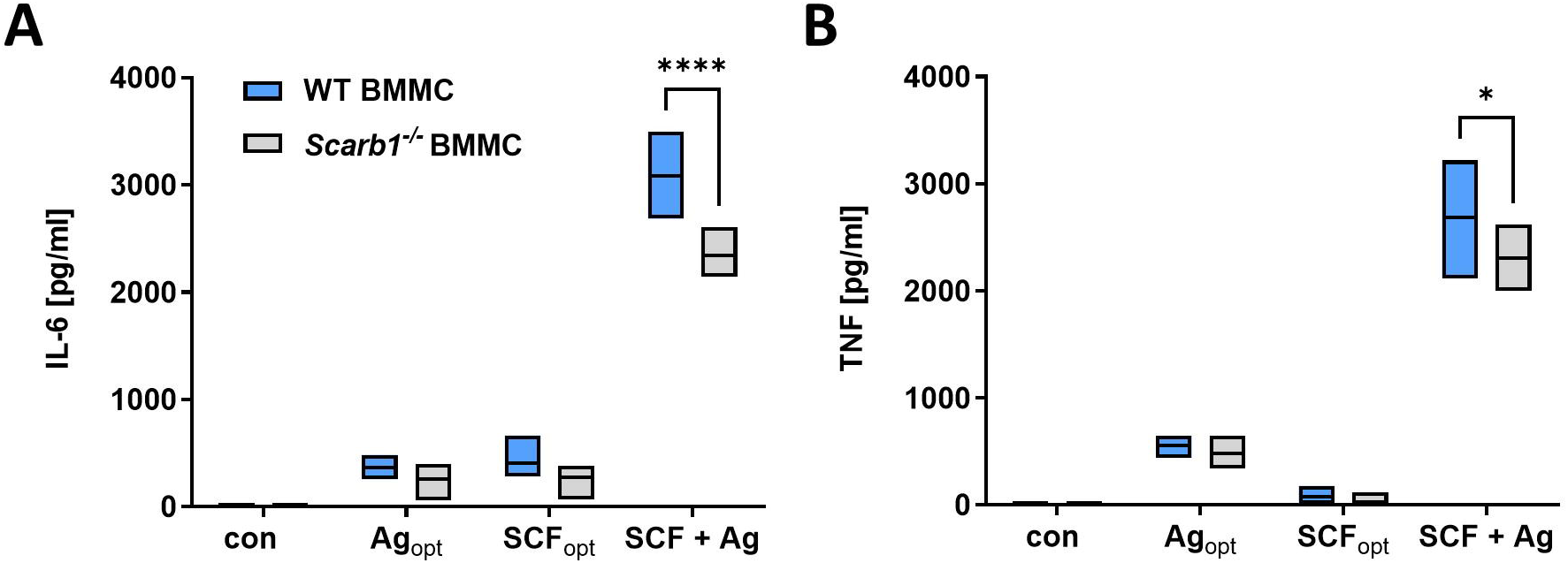
The synergistically enhanced cytokine production of BMMCs upon SCF/Ag co-stimulation is reduced in *Scarb1*-deficient BMMCs. WT and *Scarb1^-/-^* BMMCs were stimulated with Ag [20 ng/ml], SCF [100 ng/ml] or the combination of both for 3 hours. The secreted amounts of IL-6 **(A)** and TNF **(B)** were determined by ELISA. Data are expressed as mean +SD or as floating bars indicating minimal, maximal and mean values. Ordinary two-way ANOVA followed by Sídák multiple comparisons test. * *p*<0.05, **** *p*<0.0001.

**Supplementary figure 5:**
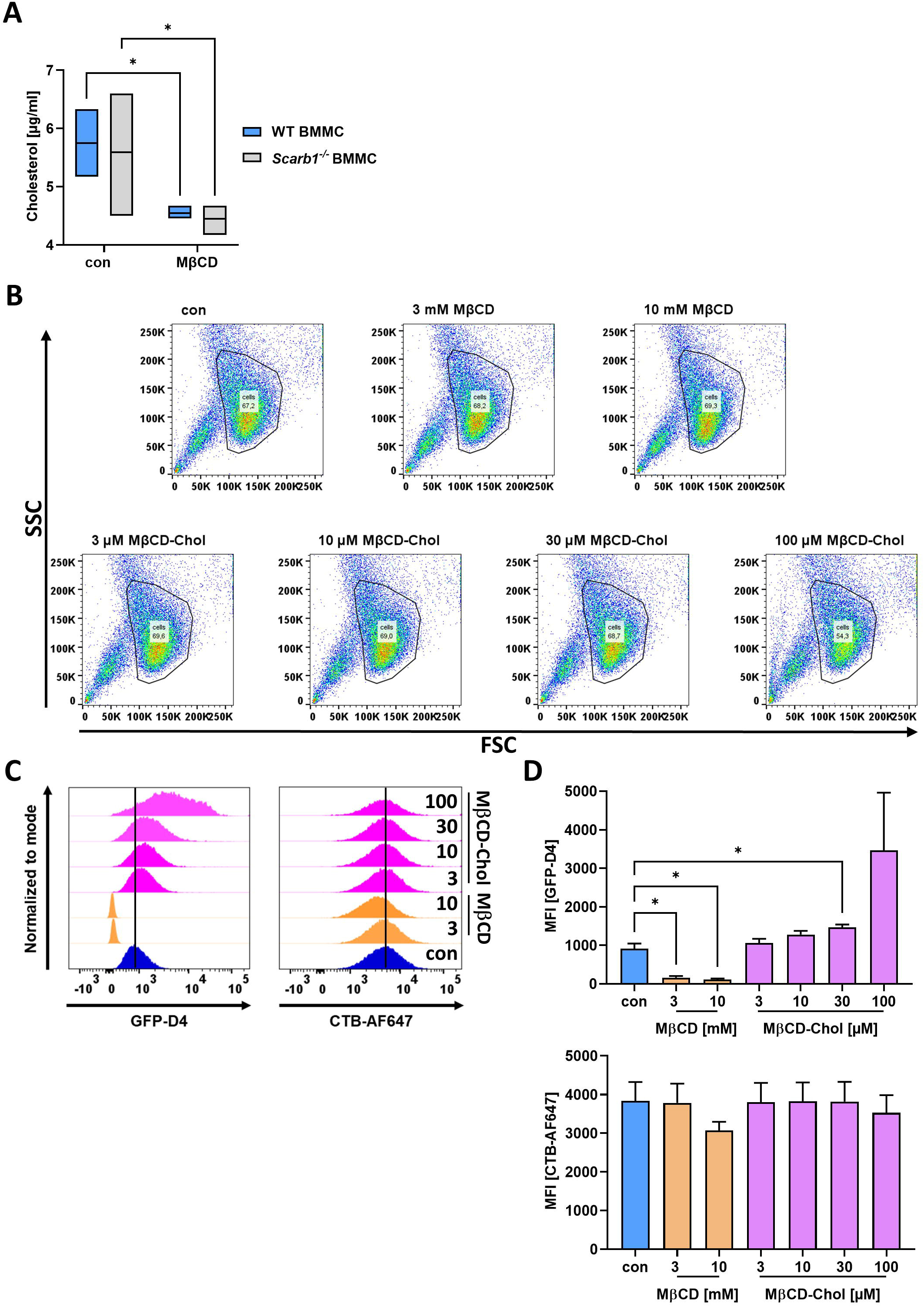
Analysis of total cellular cholesterol and plasma membrane raft associated cholesterol upon MβCD treatment. **(A)** Unesterified and esterified cellular cholesterol was measured in untreated or MβCD-treated [5 mM, 10 minutes] WT and *Scarb1*-deficient BMMCs (n=4). **(B)** Representative FACS dot plots of WT BMMCs treated with indicated concentrations of MβCD for 10 minutes or MβCD-chol for 30 minutes. **(C)** Representative histograms of WT BMMCs stained with GFP-D4 and CTB-AF647 after treatment as described in **(B)**. **(D)** Quantification of data depicted in **(C)** (n=3). Data are expressed as mean + SD or as floating bars indicating minimal, maximal and mean values. **(A)** was analyzed by two-way ANOVA followed by Sídák multiple comparisons test. **(D)** was analyzed by ordinary one-way ANOVA followed by Dunnett test to correct for multiple comparisons. * *p*<0.05.

**Supplementary figure 6:**
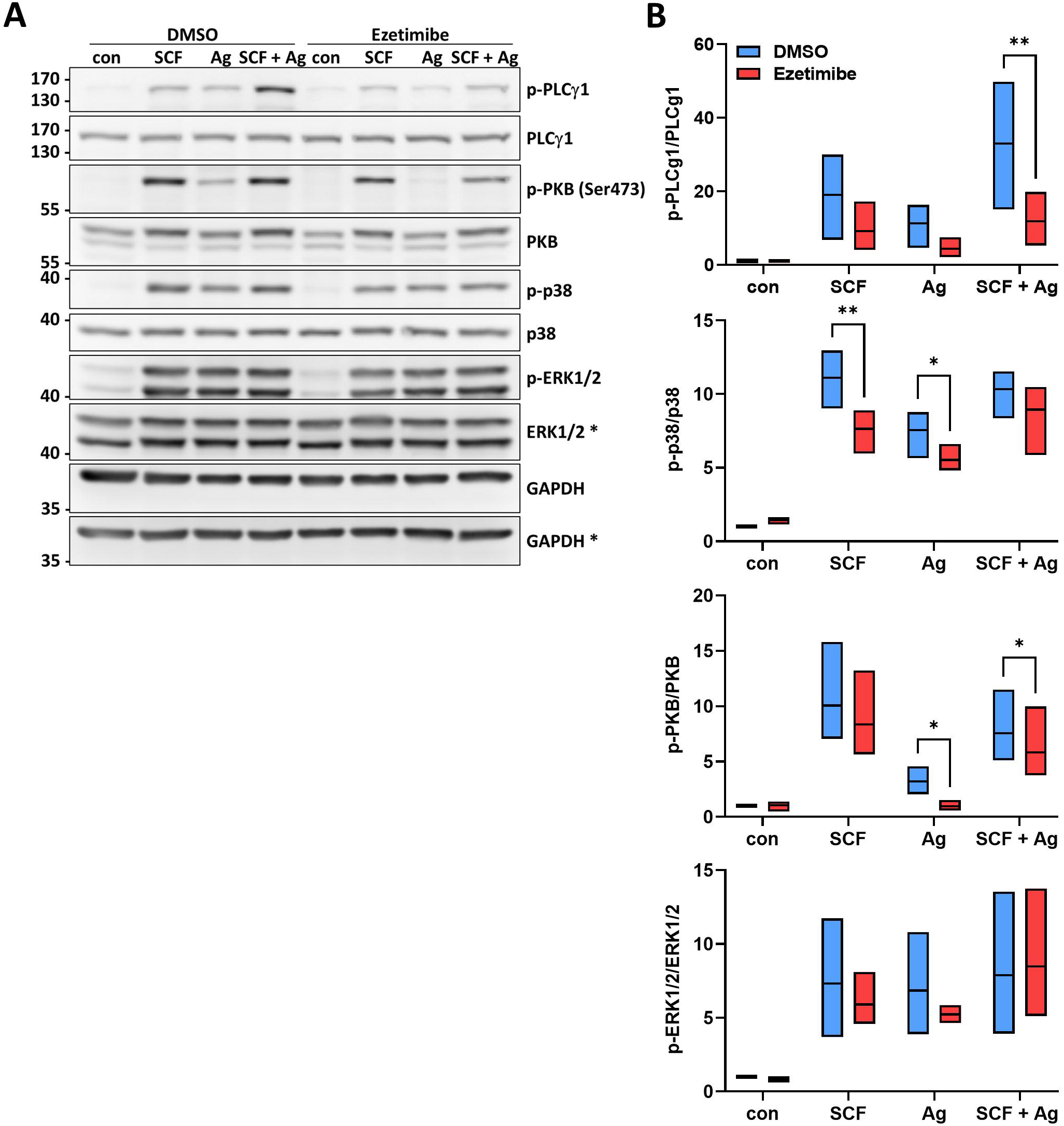
Ezetimibe reduces synergistically enhanced PLCγ1 phosphorylation in response to SCF/Ag co-stimulation. **(A)** Representative Western blot showing WT BMMCs pre-treated with DMSO or Ezetimibe [30 µM] for 1 hour and then stimulated with SCF [10 ng/ml, 10 min], Ag [20 ng/ml, 5 min] or the combination added consecutively with 5 minutes of SCF pre-stimulation. Activation of PLCγ1, PKB and MAPK pathways ERK1/2 and p38 was analyzed by phospho-specific antibodies. **(B)** Densitometry analysis of p-PLCγ1, p-p38 and p-PKB in relation to total protein amounts (n=3). Data are expressed as floating bars indicating minimal, maximal and mean values. Ordinary two-way ANOVA followed by Sídák multiple comparisons test. * *p*<0.05, ** *p*<0.01.

